# Determinants of spliceostatin reactivity at a spliceosomal zinc finger

**DOI:** 10.1101/2025.09.17.676740

**Authors:** Riccardo Rozza, Angela Parise, Jana Aupič, Angelo Spinello, Vlad Pena, Alessandra Magistrato

**Affiliations:** CNR - Istituto Officina dei Materiali (IOM) c/o SISSA via Bonomea 265, 34136, Trieste, Italy; Department of Biological, Chemical and Pharmaceutical Sciences and Technologies University of Palermo 90128 Palermo, Italy; Research Group Mechanisms and Regulation of RNA Splicing, The Institute of Cancer Research 237 Fulham Road, London, SW3 6JB

**Keywords:** RNA Splicing, Covalent Inhibition, Molecular Dynamics, QM/MM, Reaction Mechanism, Cancer, Spliceostatin A

## Abstract

Branch-site recognition is a pivotal event in spliceosome assembly, and branch-site antagonists disrupt this process, inducing a splicing rewiring that underlies much of their antitumor activity. Antagonists from the spliceostatin family covalently bind the SF3b complex nearby the branch-site pocket, forming an adduct with Cys26 of the PHF5A zinc finger while remarkably preserving Zn^2+^ coordination - an unconventional mode of covalent inhibition.

Yet the molecular features that activate this normally protected ZnCys_4_ cysteine and initiate spliceostatin’s epoxide ring opening have remained obscure. Here, we combine extensive classical and QM/MM molecular dynamics simulations to map the full reactive trajectory of spliceostatin attachment to PHF5A. We identify two alternative noncovalent spliceostatin binding poses and show that only its thermodynamically favoured conformation pre-organizes the epoxide for nucleophilic attack. The local distortion of the ZnCys_4_ coordination sphere weakens the Zn–Cys26 bond, enabling water–Cys26 exchange and generating a highly reactive nucleophilic thiolate. Covalent bond formation is subsequently accelerated by an Asp34–Lys29 proton relay that activates the epoxide leaving group. Free-energy calculations confirm that the overall reaction is fast and strongly exergonic. Together, these findings provide a complete mechanistic framework for zinc-assisted inhibition of early spliceosomes by branch-site antagonists and advance our fundamental understanding of zinc finger reactivity.

## 1. Introduction

Removal of introns and ligation of exons in pre-mRNA transcripts, i.e., pre-mRNA splicing, is an essential step of eukaryotic gene expression and diversification.^1,2^ Pre-mRNA splicing leads to the formation of protein-coding mRNA or long-noncoding RNAs. Splicing is catalyzed by the spliceosome, an intricate biomolecular engine comprising hundreds of proteins and five small nuclear ribonucleoproteins (snRNPs), U1, U2, U4, U5, and U6.^3,4^ The accuracy and precision with which splicing occurs are key for correct gene expression, as splicing errors can produce dysfunctional proteins or even disease states. Recognition of the reactive groups for the catalytic process is vital for splicing fidelity.^5,6^ The spliceosome assembles de novo at each splicing cycle and proceeds through landmark intermediate complexes (including E, A, B, C, P, and ILS). The early spliceosome complexes, E and A, are involved in recognizing and chaperoning key sequence motifs: the 5’splice site (5’SS), the branch sequence (BS), the polypyrimidine tract (PPT), and the 3’SS. Recognition of these sequences often occurs in different ways, giving rise to alternative splicing.^7^ While the 5’SS is selected by the U1 snRNP (E complex), the BS and PPT-3’SS are recognized in a cooperative manner by the SF1 protein and the U2AF heterodimer. During A complex formation, U2AF recruits U2 snRNP, which binds to the BS. The BS contains the conserved branch-site adenosine (BS-A) that acts as a nucleophile of the first splicing step.^5^

The BS selection is mediated by the heptameric SF3b complex (Figure 1A), which is part of U2 snRNP, and is further composed of SF3B1, SF3B3, SF3B5 and PHF5A proteins. Among these, SF3B1 and PHF5A form a pocket (the branch-site pocket) that accommodates the BS-A during the selection.

**Figure 1:**
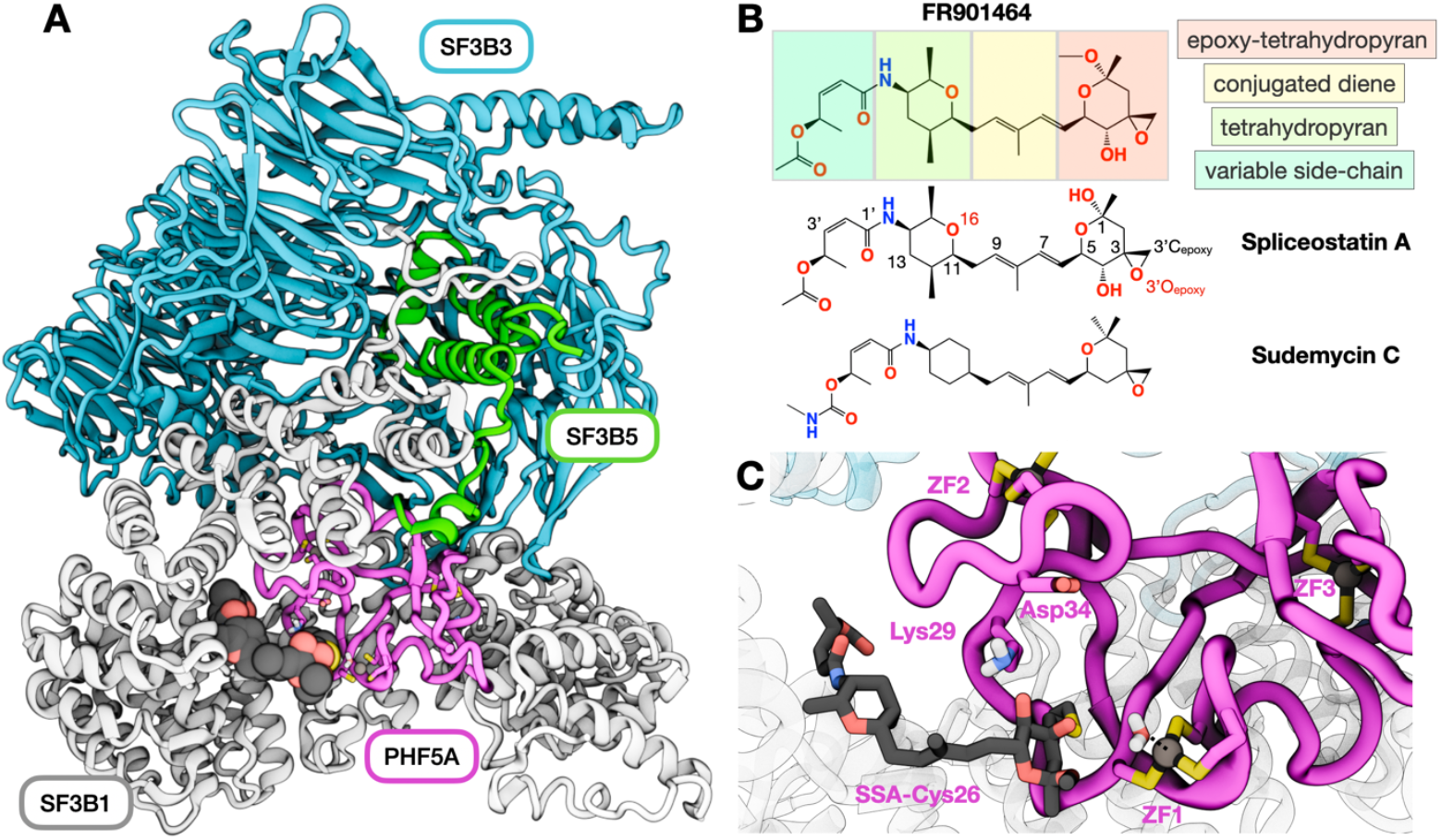
Representation of the SF3b complex and covalent inhibitors. **A** Most representative structure extracted from the 2 μs-long Molecular Dynamics simulation trajectory of the SF3b complex (based on PDB id 7b9c) in complex with Spliceostatin A (SSA). The SSA is shown in sticks. **B** Chemical structure of FR901464 and two of its derivatives, SSA and sudemycin 6 (SD6). FR901464 is made up of four parts: (i) an epoxy-tetrahydropyran, (ii) a conjugated diene, (iii) a tetrahydropyran, and (iv) a variable side chain. **C** Close-up on the PHF5A reactive site. The three zinc fingers (ZF1, ZF2, ZF3) are shown in sticks with the carbon atoms colored according to the protein to which they belong, and the zinc atoms are represented as a grey van der Waals sphere. The Cys26-SSA residue, the other cysteine residues forming the PHF5A zinc fingers, the coordinated water molecule, and Lys29 and Asp34 are shown as sticks. Carbon atoms of SSA and PHF5A are shown in grey and pink, respectively. Oxygen, nitrogen, and sulfur are shown in red, blue, and yellow colors.

Interestingly, SF3B1 and PHF5A are also upregulated in cancers.^8,9^ Most notably, SF3B1 is recognized as an oncogenic driver,^10^ that carries recurrent mutations in patient samples affected by different types of cancers, i.e. myelodysplastic syndrome with ringed sideroblasts, leukemia, uveal melanoma, and pancreatic cancer. These mutations induce the selection of cryptic BS and 3’SS instead of the canonical ones, leading to aberrant splicing events.^8,9,11,12^

SF3B1 is also a target of several splicing modulators that act as competitive antagonists of the BS-A. While SF3B1 undergoes a conformational transition from an open to closed state upon pre-mRNA binding, these splicing modulators have been shown to lock SF3B1 in the open conformation. They thus prevent BS binding^3,13–15^ and trigger intron retention or exon skipping, depending on the selected competing BS sequence and the dose of the compound. While some splicing modulators entered clinical trials,^16,17^ their clinical potential in cancer is still insufficiently explored.

Most splicing modulators targeting SF3b share a common central diene motif. Depending on the flanking chemical groups, they can be divided into three distinct chemotypes: herboxidiene, pladienolides, and spliceostatins (the latter belonging to the FR901464 family, Figure 1B).^11,12^ All three families contain epoxy groups – highly reactive electrophiles that can interact covalently with nucleophilic groups. However, crystal structures have revealed that only spliceostatins, and the related sudemycins, can employ the epoxy group for covalent coupling.^18–22^ Indeed, a recent crystal structure of two FR901464 analogs, spliceostatin A (SSA) and sudemycin 6 (Figure 1B), revealed that these modulators outcompete the BS-A and stall the spliceosome by covalently linking to a cysteine (Cys26) residue of one PHF5A zinc finger. Notably, the PHF5A residue Lys29 was suggested to contribute to coupling efficacy, given its proximity to the SSA epoxide and the observed change of sensitivity to SSA in cells carrying K29A mutations.^23^ Besides rationalizing the biological activity of the spliceostatin family of compounds, this finding demonstrated that splicing modulation could be achieved via an unconventional mode of covalent inhibition.

Covalent inhibition generally occurs through a series of discrete steps (Scheme S1): initially, the inhibitor (I) diffuses through the solution to encounter the target protein (E). Once bound, the non-covalently associated inhibitor undergoes conformational adjustments within the binding pocket (E·I) until the electrophilic warhead is optimally positioned to react with the nucleophilic residue of the biomolecular target. Next, the chemical reaction occurs, ultimately leading to the formation of a covalent bond between the inhibitor and the biomolecule (E-I).^24^

Numerous covalent inhibitors target cysteine residues and their mechanism of binding is well established.^25–27^ However, covalent inhibitors targeting zinc-associated cysteine ligands are rare. While the prevalence of zinc fingers in proteins makes this motif an attractive target for future drug design campaigns, successful design of covalent inhibitors demands a precise understanding of each step of their binding process.

Here, we performed classical and hybrid quantum-classical (QM/MM) molecular dynamics (MD) simulations to elucidate the molecular mechanisms that usher the covalent coupling of SSA to SF3b. We elucidate how SSA binds PHF5A in a reactive site of SF3b, reminiscent of an enzymatic center microenvironment. Namely, Cys26 responsible for the nucleophilic attack on the SSA warhead originates from and is activated by a distorted zinc finger of the PHF5A protein. Additionally, the proton relay of Asp34 and Lys29 pair of PHF5A residues assists the SSA covalent linkage by protonating the dissociating oxygen atom of the warhead (Figure 1C). These findings advance our fundamental understanding of zinc finger reactivity and could aid the development of novel covalent inhibitors targeting zinc finger motifs in spliceosome proteins and other biological contexts.

## 2. Results

### 2.1. SF3b stabilizes a reactive L-shaped SSA conformation and alters Zinc Finger in the pre-reactive complex

To determine whether SF3b preorganizes SSA into a reactive conformation and perturbs the target zinc-finger prior to covalent bond formation, we simulated the conformational dynamics of SSA and the geometry of the reactive cysteine in the non-covalent E·I complex.

To relax the SSA/SF3b non-covalent adduct (derived according to the procedure in Method Section 4.1) and to describe the set of non-covalent interactions that SSA establishes with SF3b we perform multi-replica 2 μs-long classical MD simulations with the FF14SB^28^ AMBER FF (Figure S1). Here, the SSA sidechain and tetrahydropyran moiety (Figure 1B) hydrogen bonded with residues Arg1074 and Leu1066 from SF3B1 and formed hydrophobic interactions with Lys1067, Val1078, Val1110, Val1114, and Tyr1157 from SF3B1 and Lys25, Tyr36, and Val37 from PHF5A (Figure S1, Table S1).

Interestingly, SSA displayed substantial conformational variability, transiting multiple times during the simulation between a bent L-shape, trapped in the X-ray structure, and a linear I-shaped conformation (Figure 2). This transition occurred via rotation around the bond linking the tetrahydropyran ring to the conjugated diene (dihedral angle ω_O16,C11,C10,C9,_ Figure 2). The ω_O16,C11,C10,C9_ dihedral angle value was ∼-180° and ∼ -60°, for the L and I-shape states, respectively (Figure 2, S2). This rotation affected the position of the epoxy ring warhead, which lied at a distance of 22.1 and 5.9 Å from Cys26-S in the I-shape and L-shape conformations, respectively. Conversely, the SSA sidechain (Figure 1B) remained anchored to SF3B1 by hydrogen-bonding to Arg1074 and Leu1066, and the SSA diene moiety was stabilized by Tyr36-PHF5A hydrophobic interaction (Figure S1). We next computed the free energy profile (Figure 2B) associated with the SSA conformational transition by analyzing the probability distribution of the I- and L-shape states during the MD simulation. We found that the L-shape state is the most thermodynamically favorable (ΔG= -1.9 ± 0.4 Kcal/mol) and that the free energy barrier (ΔG^‡^) associated with L to I-shape interconversion is 3.1 ± 0.4 Kcal/mol. This corresponds to a conversion rate of k=1.5±1.0 ·10^10^ s^-1^ as estimated via the Eyring–Polanyi equation. The SF3b complex is responsible for the larger stability of the reactive L-shape SSA conformation. Indeed, metadynamics simulations (details in Method Section 4.2.3) of SSA in water revealed that the SSA I- and L-shape conformations are thermodynamically equivalent with a ΔG= 0.3±0.2 kcal/mol (Figures 2C,D). In water solution, a further (bent-shape) minimum at ω_O16,C11,C10,C9_=+46.8° was discovered, which was not accessible in the protein environment due to steric hindrance (Figures 2C,D). However, SF3b did not affected the I- and L-shape interconversion rate, since in water SSA exhibited a similar free energy barrier (ΔG^‡^ =3.4±0.2 kcal/mol) to that computed in presence of SF3b. Quantum Mechanical (DFT-B3LYP) calculations in implicit solvent further confirmed these results (Figures S3). As such, SSA might explore distinct conformational states before anchoring to Arg1074 and Leu1066 residues and adopting a reactive conformation.

**Figure 2:**
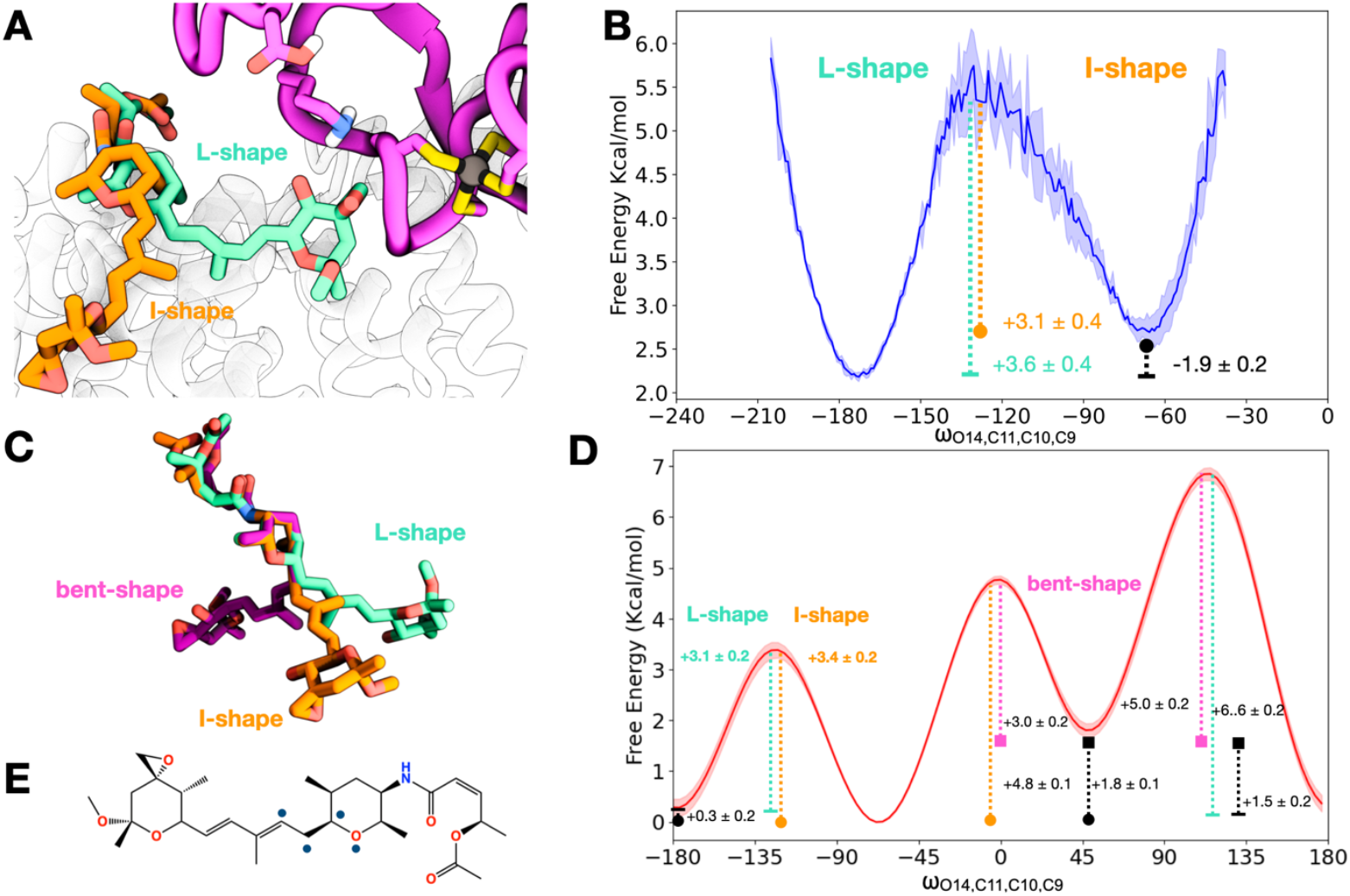
Conformational Analysis of SSA. **A** The two SSA conformations achieved by rotation along the ω_O14,C9,C10,C11_ dihedral angle of spliceostatin A in the SF3b complex. The I-shape and L-shape SSA carbon atoms are depicted in light mint and orange, respectively. **B** Free energy profile (Kcal/mol) for rotation along the SSA ω_O14,C9,C10,C11_ dihedral angle in the SF3b complex obtained from classical molecular dynamic simulations. The standard deviation is depicted as a blue shadow. The energy level of L- and I-shape conformations are identified with a line or a circle and located at -172,8° and -65,7*, respectively. **C** The three available conformations of SSA in water and in absence of the Sf3b complex. The I-shape and L-shape SSA carbon atoms are depicted in light mint and orange, respectively. An additional bent conformation accessible only in water solution is reported in magenta. **D** Free energy profile (Kcal/mol) for the rotation of SSA along the ω_O14,C9,C10,C11_ dihedral angle in water complex obtained from classical metadynamics simulations. The standard deviation is depicted as a red shadow, obtained by averaging 3 replicas. The I-shape minimum is located at -68.4° and is set as the reference level for energy. The L-shape minimum is located at -176.4° A further minimum (black square, not explored in the presence of the Sf3b complex) was found at +46.8°. The free energy barrier to interconvert from the I to the L-shape state is 3.4±0.2 kcal/mol. **E** Schematic structure of SSA with blue dots pointing at the atoms defying the ω_O14,C9,C10,C11_ dihedral angle.

Next, we relaxed the E·I state, considering SSA in the L-shape conformation, by performing 5.5 ps of QM/MM MD simulations. We used the DFT-BLYP-D3 level of theory for the QM region, which included the reactive moieties (Figures S4A,B): the reactive zinc finger, Asp34 and Lys29 residues and the reactive part of SSA (up to C10, Figure 1B) and the same force field of the classical MD simulation for the MM part (details in Method Section 4.2.2). This method allows the description of the reactive center of a biomolecule at an accurate QM level of theory and can account for rearrangements of electronic structure occurring during chemical reactions, while the rest of the system is treated with classical force fields.^29^ Notably, the reactive zinc finger geometry departed from the canonical architecture. Indeed, Cys26 exhibited a more plastic Zn-S coordination bond than the other Zn-S bonds (Figure, 3) S(Cys-26)-Zn bond laid for 46.6% of the time above 2.45 Å, while the bond length of the remaining Zn-S-Cys bonds overcame the typical zinc-finger Zn-S coordination bond length of 2.45 Å less frequently (Zn and S-Cys23, -Cys58, and -Cys6 41.3%, 30.9%, and 40.5% of the simulation time, respectively). Concomitantly, the zinc finger also deviated from the ideal tetrahedral geometry. Indeed, out of the six S-Zn-S angles, only three exhibited interquartile ranges encompassing the ideal tetrahedral value of 109.5° (Figure 3).

**Figure 3.**
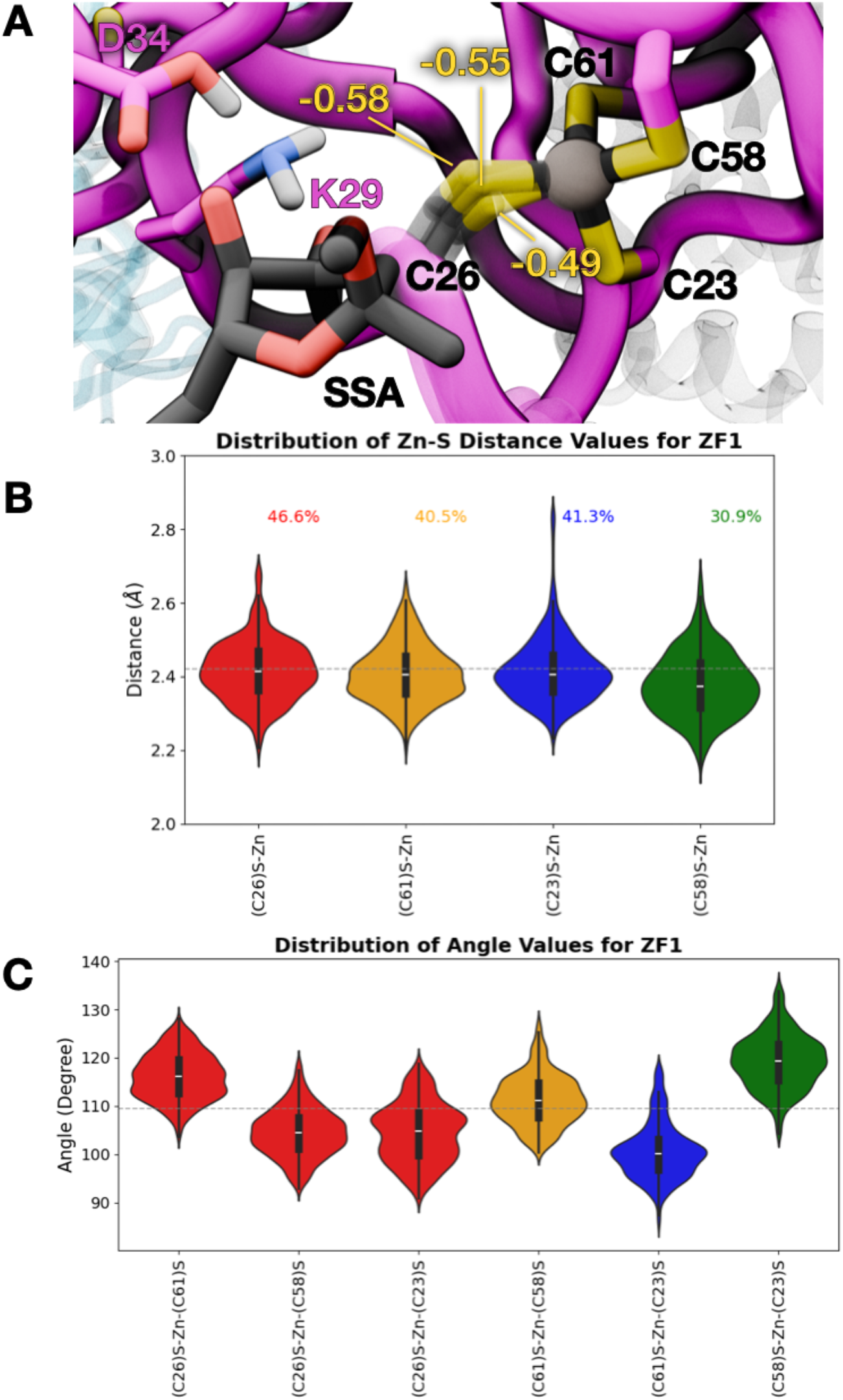
Distortion of Zinc Finger 1: (**A**) Close view of SSA binding site showing the distortion of Zinc Finger 1. Fluctuations of Zinc-S-Cys26 dsitances sampled during QM/MM MD simulation of the E·I state. Computed partial changes (e) of Cys26 sulphur atom corresponding to each distance are shown in yellow (**B**) Violin plots reporting the distributions of Zinc-sulphur distances (Å). The dashed line, at 2.42 Å, demarks the limit of the of the Zn-S coordination bond length. The percentage of QM/MM MD trajectory frames where this limit is overcome is shown on top of each dataset; The distribution of the reactive Zn-S(C26) distance is coloured in red, those of the Zn-S(C61), Zn-S(C58), Zn-S(C23) are depicted in yellow, blue and green, respectively. (**C**) Violin plots reporting the distributions of the Sulphur-Zinc-Sulphur angles (degree). Angles involving the reactive Cys26 residue are coloured in red, those of (C61)-S-(C58)S, (C61)-S-(C23)S, and (C58)-S-(C23)S are coloured in orange, blue, and green, respectively. The dashed line at 109.5°demarks the value an ideal tetrahedral coordination sphere.

As such, our simulations reveal that SF3b thermodynamically favors the L-shaped reactive conformation of SSA relative to the I-shaped state and promotes increased structural plasticity of the zinc-finger cysteine in the pre-reactive E·I complex, thereby favoring a geometry competent for covalent crosslinking.

### 2.2. Covalent inhibition mechanism of spliceostatin A

How spliceostatin A converts a pre-reactive binding pose into an irreversible covalent adduct within the SF3b complex remains unclear. To address this, we dissected the atomistic reaction pathway and free-energy landscape of SSA covalent binding starting from the non-covalent E·I state, previously equilibrated via QM/MM MD simulations (Figure 4 and Section 4.2.5).

**Figure 4.**
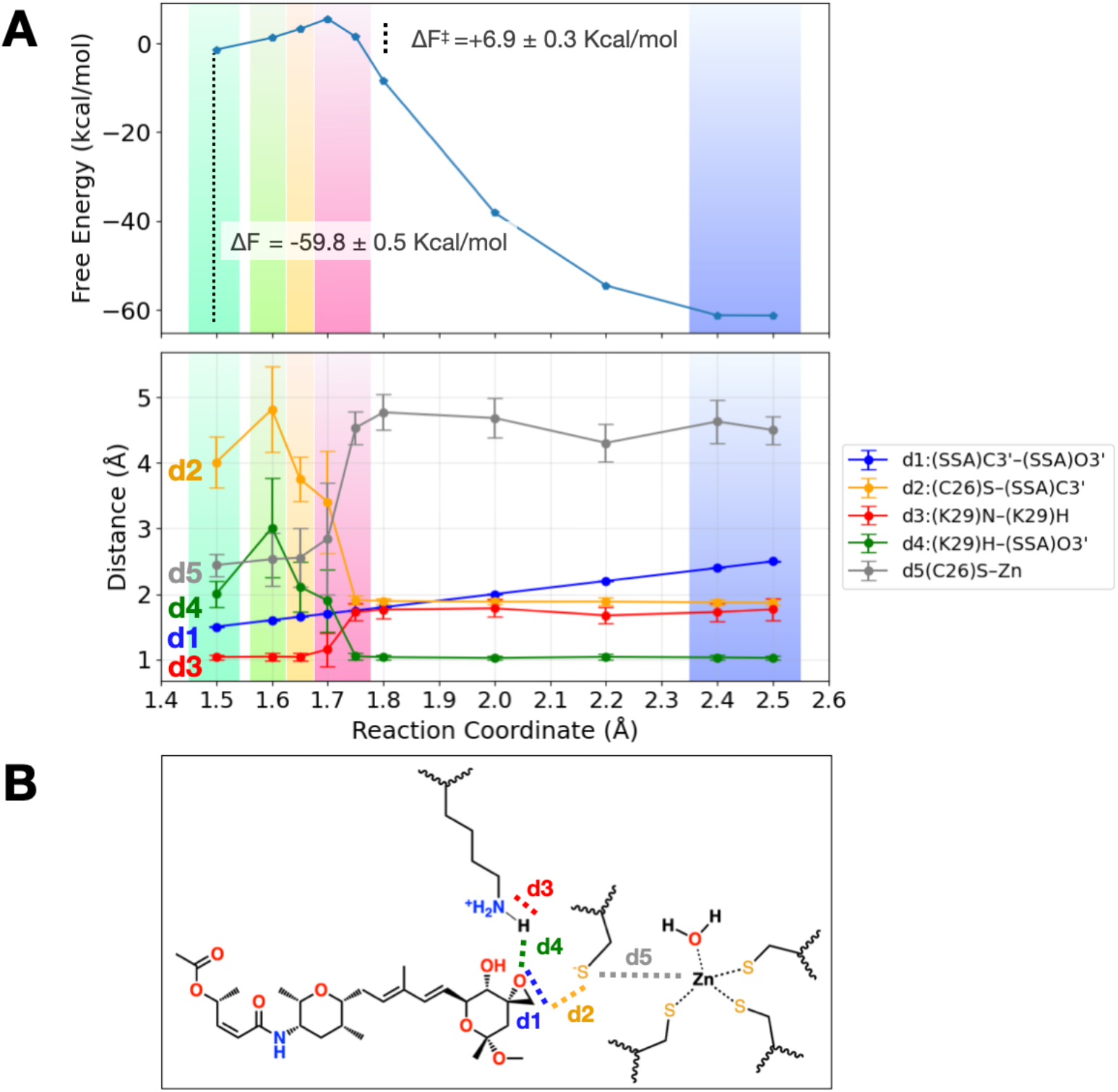
Energetics of spliceostatin A covalent binding. (A) Free Energy profile (kcal/mol) for the covalent linkage of spliceostatin A to PHF5A (top) and mean values of selected distances between reactive atom as a function of the reaction coordinate (bottom). (B) Schematic representation of the distances monitored along the reaction coordinate.

We next applied biased QM/MM MD simulations combined with the thermodynamic integration method and using as reaction coordinate (RC) the distance between the epoxy carbon and oxygen atoms (3’C and 3’O, Figures 1 and S4A) of SSA. We performed QM/MM MD simulations on RC windows ranging from 1.5 Å to 2.4 Å. The RC was chosen basing on QM/MM SteeredMD simulation (Details in Method Section 4.1)

In the reactant state (RC=1.50 Å), the Cys26-S coordinated the Zn ion, and the 3’O-epoxy atom of SSA was locked into a reactive conformation by an extended interaction network involving Lys29-Nζ-PHF5A and Asp34-Oγ-H (Figure 5 and Scheme 1 in Section 3). Proceeding along the RC (1.60Å), the Asp34-Oγ-H proton was released to Lys29, which becomes ionized (Scheme 1).

**Figure 5.**
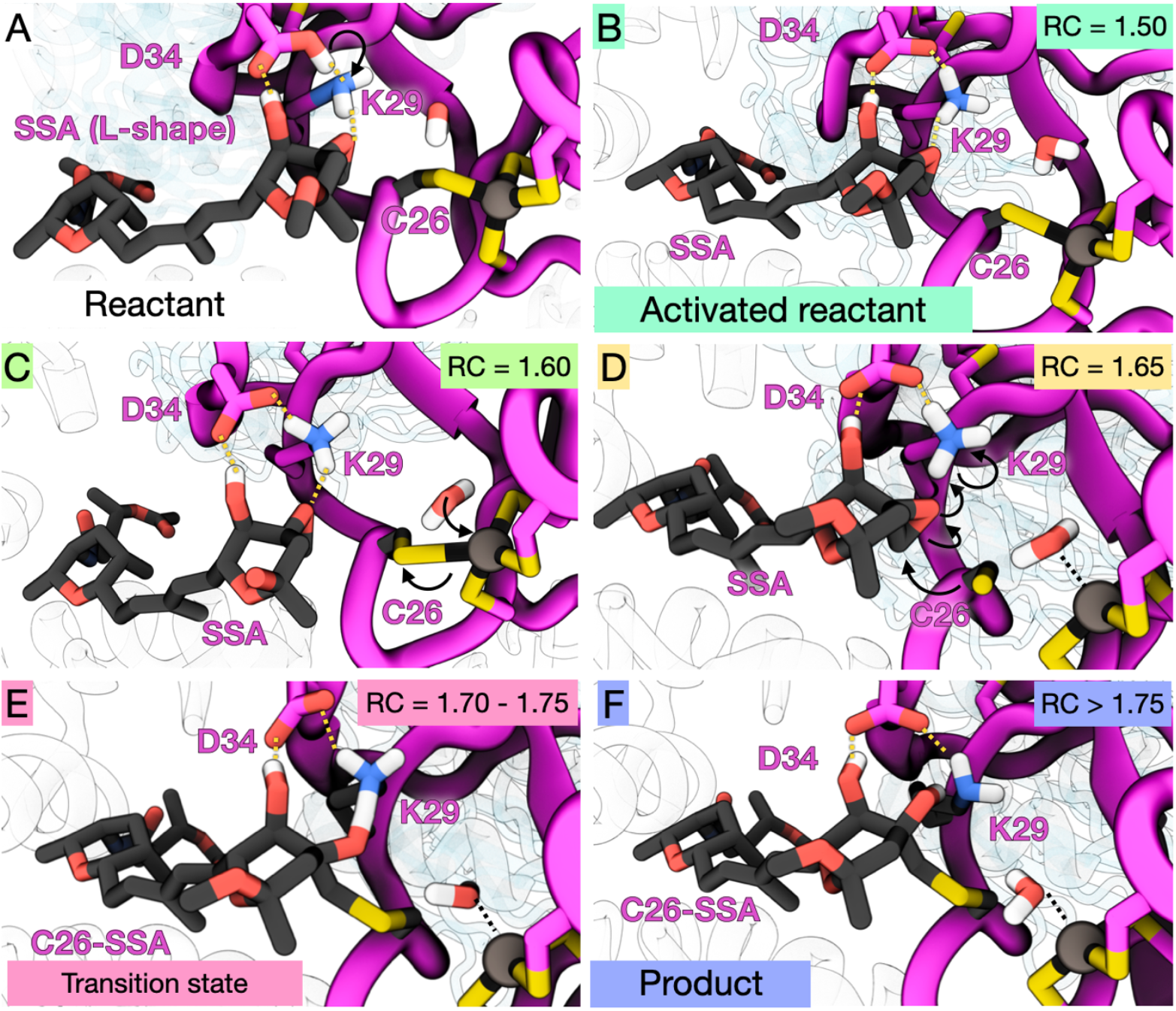
Molecular mechanism of SSA covalent binding to PHF5A. (**A**) Pre-reactive state: SSA in L-shape configuration approaches K29, D34 and C26. (**B**) Activated Reactant (RC=1.5Å, light mint), D34-H protonates K29. (**C**) Nucleophile activation (R=1.60Å, lime green), a water molecule enters the zinc coordination sphere and (**D**) (RC=1.65Å, pastel yellow) the C26-S-Zn coordination bond breaks and becomes ready to perform the nucleophilic attack, triggering epoxide ring opening and protonation by K29. Black arrows show electron movements proceeding along the reaction coordinate. (**E**) Transition state (RC=1.70-1.75 Å, peach pink) featuring the nucleophilic attack of C26-S atom on the C3’ carbon atom of the SSA-epoxy ring, C3’-O3’ epoxy bond breakage and, the SSA-O3’ protonation by K29. (**F**) Product state (RC=2.4-2.5 Å, cornflower blue). Only the hydrogen atoms involved in the reaction or in the interaction with PHF5A are shown.

At RC=1.60-1.65 Å, a water molecule approaches the coordination sphere of the zinc ion, triggering Cys26-S dissociation and priming it for the nucleophilic attack on the 3’C-epoxy atom of SSA. Here a water molecule substitutes the Cys26-S ligand. At RC=1.70-1.75 Å, the transition state is reached, and the free energy profile reaches its maximum, corresponding to a free energy barrier of ΔF^‡^= 6.9 ± 0.3 Kcal/mol (Figure 4) and a reaction rate of k=8.8±4.3 ·10^7^ s^-1^, as estimated by the Eyring–Polanyi equation. At RC=1.70, the 3’O-epoxy and 3’C-epoxy bond starts breaking due to the formation of the 3’C-epoxy and Cys26-S covalent linkage. At RC 1.75 Å, the 3’O-epoxy abstracts a proton from Lys29-Nζ, completing the covalent binding process. For higher RC values (2.0 – 2.4 Å), the SSA epoxy ring warhead is fully transformed into a hydroxyl group, which hydrogen bonds to Lys29-Nζ, and the C3’-S-Cys bond between SSA and the PHF5A protein is formed (Figure 4). Overall, the nucleophilic substitution reaction between Cys26-S and the O3’-epoxy at C3’ of SSA follows an SN2-type mechanism. Here, the breakage of the 3’O-epoxy and 3’C-epoxy bond is endorsed by the S-Cys nucleophilic attack and supported by the proton transfer from Lys26-Nζ (Scheme 1, Figure 5, Figure S5, Figure S6). Importantly, Asp34 orients and acidifies Lys29, thus supporting its role as a general base. This process is highly exergonic with ΔF=-59.8±0.5 Kcal/mol (Figure 4) consistently with its irreversibility.^30^ 7 ps of QM/MM MD simulations of the product state revealed that 3 Zn-Cys bonds, a water molecule form the coordination sphere of the zinc ion. Subsequent 2 μs long MD simulations confirmed that SSA stably bound to SF3b (Figure S7), decreasing its flexibility (Figure S9-S12). Overall, these simulations revealed a stepwise SN2-type mechanism in which zinc coordination remodeling, water-assisted Cys26 activation, and Lys29/Asp34-mediated proton transfers cooperatively enable epoxide ring opening and covalent bond formation. The low free-energy barrier and strongly exergonic profile explain the efficiency and irreversibility of SSA covalent linkage to PHF5A.

## 3. Discussion

Small-molecule covalent drugs have several advantages in comparison to their non-covalent counterpart. Most prominently, the formation of a covalent bond between an inhibitor’s reactive functional group and its target protein results in enhanced binding affinity that significantly surpasses that of non-covalent interactions alone. The covalent bond leads to an extended retention time of the inhibitor, contributing to unique pharmacodynamic characteristics that yield remarkable potency. The high efficacy of covalent inhibitors can occasionally lead to toxicity issues.^25,31,32^ While first covalent drugs were discovered serendipitously and often lacked selectivity, covalent drug design has more recently yielded several successes,^24,25^ sparking a renowned interest in this field. Design of efficient covalent inhibitors requires a detailed understanding of the configuration of the reactive centers and how these facilitate the reaction process, yet mechanistic information on their covalent binding is most often lacking. Computer simulations can complement X-ray crystallography and cryo-EM structural studies with kinetic and thermodynamic insights that include conformational dynamics, identification of transient states, and free energy barriers for bond breaking and formation. Therefore, they are essential for pinpointing key chemical moieties involved in covalent bonding and for offering detailed guidance for the future development of covalent inhibitors.

Here, by performing classical and QM/MM MD simulations, we detailed the mechanism of covalent bond formation between the splicing modulator SSA and a cysteine residue of a PHF5A zinc finger. We observed that during the non-covalent association, SSA established extensive interactions with the reactive center – adjacent to the branch site pocket, at the interface of the PHF5A and SF3B1 proteins (Figure 1).^33^ However, SSA was firmly bound only at its side chain and diene moiety, while the rest of the molecule was characterized by large conformational variability. Indeed, SSA established assumes two conformations: an I-shape and a reactive L-shape, which is thermodynamically more stable, thanks to a specific set of interactions established with the SF3b. In the L-shape state, the SSA epoxy moiety was optimally oriented to undergo a nucleophilic attack by one cysteine of a PHF5A zinc finger. Notably, the zinc finger ion takes an active role in this process by facilitating the conversion of the coordinated cysteine into an anionic nucleophilic thiol group. The geometric restraints imposed by the PHF5A protein divert this zinc finger from a canonical tetrahedral geometry (Figure 3). This distortion is reflected in the frequent elongation of the reactive Cys26 ligand, making it susceptible to dissociation from the zinc ion through a water-ligand exchange reaction. As such, Cys26 becomes readily available to perform a nucleophilic attack on the epoxy warhead of SSA. An electronic structure analysis of the reactive zinc finger revealed that the sulfur atom of the most frequently elongated Cys26 possesses a larger negative charge as compared to the sulfur atoms of the remaining cysteines. This is in line with its weaker coordination to the zinc ion (Figure 3 and S13), and its enhanced nucleophilic character. Consistently with this finding, an atomic force microscopy revealed that zinc fingers containing four cysteines are characterized by high kinetic lability.^34,35^ Therefore, this type of zinc fingers is particularly inclined to Zn-S bond dissociation and amenable to act as nucleophilic agents. Specifically, Cys26 is part of a zinc ribbon-type motif in which each couple of cysteine ligands are contributed by two different knuckle and lie in close proximity, being separated by only two-residues.^36^ The tension imposed by this structural arrangement may be responsible for the observed zinc finger distortion, resulting into the elongation, enhanced flexibility and reactivity of Cys26.

In addition to the role of the zinc ions in promoting nucleophile activation, the PHF5A protein actively promotes SSA covalent binding by exposing a unique acid-base residue pair in proximity to its epoxy moiety. Reminiscent of a domino effect, the nucleophilic attack of Cys26-S on the SSA epoxy group stems from the donation of a proton from Asp34 to Lys29, which subsequently releases its proton to the oxygen atom of the SSA epoxy group (i.e., the leaving group in the nucleophilic substitution reaction). This facilitates the O3’ epoxy dissociation from the C3’ atom and its conversion into a hydroxyl group. The formation of the covalent linkage between SSA and PHF5A is highly exoergonic (ΔF= -59.8 ± 0.5 Kcal/mol) in line with an irreversible inhibition process.

**Scheme 1:**
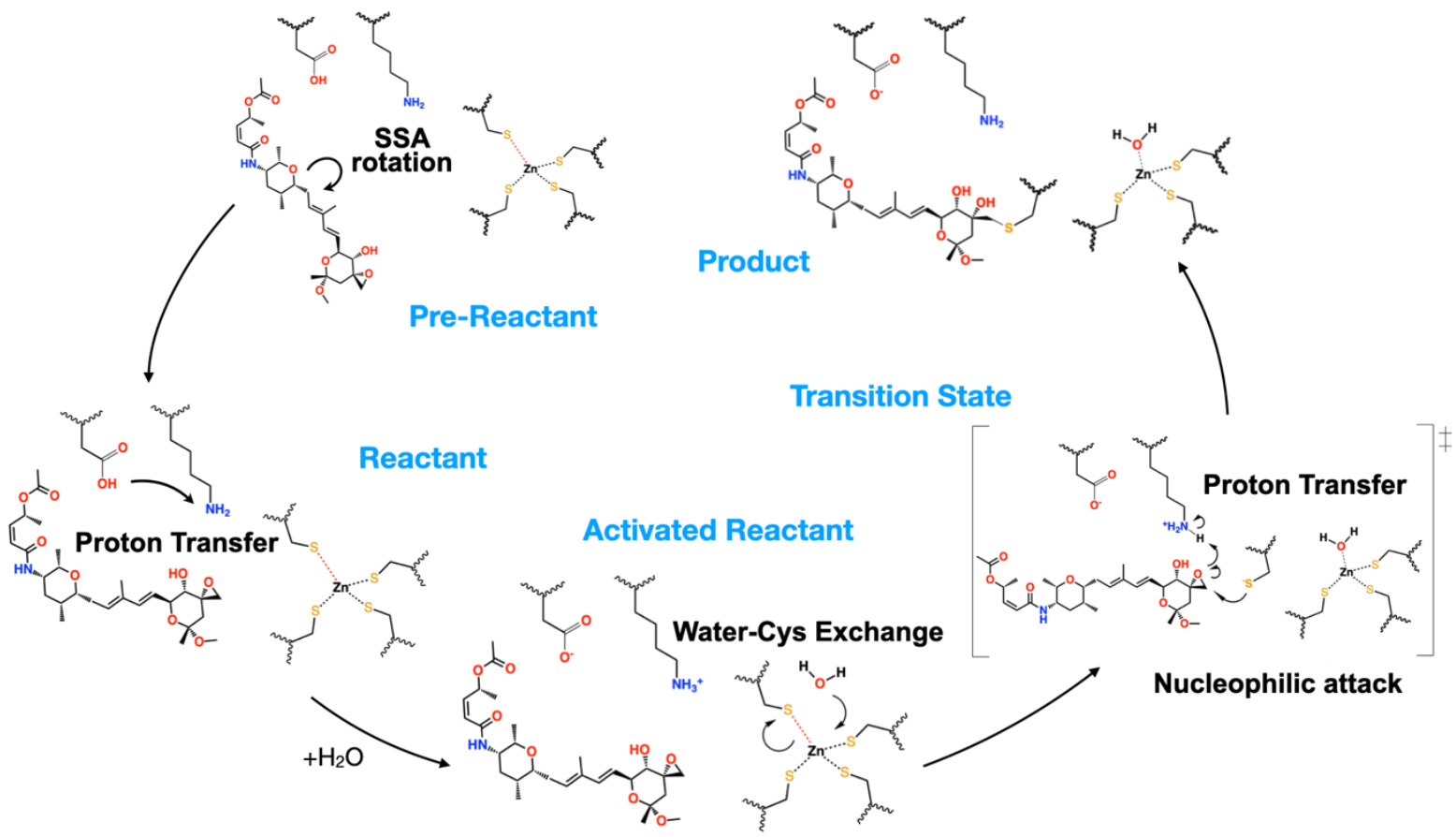
Schematic picture of the reaction Mechanism of SSA covalent binding to PHF5A. The distorted Sulfur-Zinc bond of Cys26 is highlighted with a red dashed line. In pre-reactant sate, SSA transit from an I-to-L shape conformation upon rotation of the dihedral angle. In activated reactant state, Asp34 protonates Lys29 and a water molecule triggers Cys26 dissociation from the zinc coordination sphere, by approaching the zinc ion. In the transition state, Cys26 performs a nucleophilic attack on the SSA epoxy carbon atom and the SSA epoxy oxygen atom extracts a proton from Lys29. In the end, the product is formed, and the newly generated hydroxyl group is stabilized by hydrogen bonding to Lys29.

The observed mechanism (Scheme 1), likely shared by other FR901464 family of drugs, has the chemical features reminiscent of a metal-assisted enzymatic process: (i) the residues of the binding site specifically recognize the drug and stabilize it into a reactive (L-shape) conformation, (ii) the zinc ion assists the formation of a nucleophilic Cys moiety, (iii) the Asp34/Lys29 acid/base couple promotes the release of the leaving group. Thus, this work demonstrates that drug neighboring residues participate in endorsing the epoxy ring covalent coupling both directly – as a proton source - and indirectly by anchoring the non-reactive drug moiety. Our findings carry important implications for future drug design campaigns, suggesting that the correct positioning of the epoxy warhead relative to the zinc motif is not the only feature to consider when selecting candidates for in vitro screening. Taking into account the residues surrounding the epoxy warhead, the geometric characteristics of the zinc finger coordination sphere seem to be equally significant.

Moreover, while cysteine residues are targets of a variety of covalent drugs,^39^ activation of a nucleophilic cysteine by zinc ions is rare and has been previously observed only in cysteines belonging zinc coordination sphere of the metallo-β-lactamase enzymes.^40–42^ However, zinc finger motifs are commonly found in a wide variety of proteins where they provide structural integrity and maintain functional shapes under a range of physiological conditions. Hence, owing to their abundance and biological relevance zinc fingers are a tantalizing target for covalent drug discovery efforts. To estimate the presence and the frequency of distorted ZnCys_4_ motifs, we analyzed their characteristics in other splicing factors and splicing-related proteins as a model of the whole proteome (Figure S14, Table S3-S6). We identified several proteins containing distorted zinc finger motifs, where at least one zinc-S-Cys distance was longer than 2.4-2.8 Å (Table 1). Most of the identified zinc finger have the same structural characteristic of the reactive PHF5A zinc finger and showcase a ribbon-like zinc finger architecture (i.e. four ligands coming from two knuckles and being separated by two residues^36^ (Table S6). Moreover, in many of them a basic residue (Lys, His, Arg) is present nearby, and it is flanked by an acidic residue (Glu, His), thus resembling the acid-base pair of residues assisting the binding of SSA to SF3b (Table S6). This analysis suggests that, upon the binding of the appropriate inhibitor/warhead, the covalent linkage mechanism elucidated here for SSA may also be applicable to other zinc finger-containing proteins.

**Table 1.** List of splicing-related proteins containing zinc finger motifs (first column) and their deposited PDB ID (second column)

| Protein | PDB ID |
| --- | --- |
| RBM5 | 7PDV |
| RBM10 | 2MXV |
| RBM22 | 6ID0, 6ID1 |
| SLU7 | 7W5A |
| ZBTB7A | 7EYI |
| SF3A3 sub 3a | 7EVO, 7VPX, 8HK1 |

Nonetheless, due to their inherent kinetic lability, even canonical (undistorted) zinc finger motifs may be targeted by covalent inhibitors and therefore it may be worth considering them in drug discovery efforts.

Interestingly, Lys29 hyperacetylation, occurring under stress condition, was recently shown to induce alternative splicing, changing mRNA levels and, consequently, expression levels of several genes. These included the KDM3A gene, encoding for lysine demethylase 3A, which promotes colon carcinogenesis.^37^ These findings suggest that the microenvironment around the branch-site from SF3b can act as a reactive center that modulates splicing, either as a natural mechanism in stress response (hyperphosphorylation) or by covalent modulators (such as spliceostatins, sudemycins and tryptoline amides).^38^

In conclusion, the zinc finger motif represents an appealing underexplored target for covalent drug discovery initiatives, and our findings advance our fundamental understanding of zinc finger reactivity and provide essential insights for designing novel zinc-assisted covalent inhibitors.

## 4. Methods

### 4.1 Generation of the E·I and E-I models

The mode of the covalent E-I complex was built on the X-ray deposited structure (PDB code 7B9C).^23^ To generate instead the non-covalent E·I complex we performed steered QM/MM MD simulations (SMD). This method operates via the application of an external force on specific reaction coordinate ^43,44^ and can be also used to identify putative reaction pathways, without providing an accurate estimate of the associated free-energy profile.

We started SMD simulations from a selected frame of unbiased QMMM MD simulation of the E-I complex. To reach the E·I state, we tested different reaction coordinates (RCs), which must enforce SSA dissociation from the zinc finger (E-I state) and reform the SSA epoxy moiety. Among the RCs tested, the distance between the epoxy carbon and oxygen atoms (3’C and 3’O, Figures S4) of SSA lead to E·I restoration.

Key to regenerate E·I in SMD simulations was the protonation state of Lys29 and Asp34 residues. Indeed, the Lys29^+^/Asp34^-^ protonation state suggested by PropKa software^45^ (Table S2) for the E-I complex did not lead to formation of the E·I state. Furthermore, during QM/MM MD simulation of the product E-I state, we observed a continuous proton exchange between Lys29^+^/Asp34^-^), indicating that their protonation states are coupled (Figure S4C). Conversely, considering the Lys29/Asp34^-^ as the starting protonation state of E·I complex SMD simulations successfully generated the E·I state.

### 4.2 Computational details

#### 4.2.1 Molecular dynamics parameters

Starting from the X-ray structure of SF3b containing SF3B1 with SSA covalently bound to PHF5A Cys26 (PDB code 7B9C, resolution of 2.40 Å).^23^ We generated three different topologies, one for the non-covalent E·I complex considering the Lys29/Asp34 protonation state, and two for the covalent E-I complex: one with Lys29^+^/Asp34^-^, namely E-I’, and another one with Lys29/Asp34^-^, being the latter the results of the covalent coupling and therefore will be referred hereafter as the E-I state.

Topologies for all studied systems were generated using the tleap module of AmberTools22.^46^ The FF14SB^28^ AMBER force field (FF) was used for the protein residues. The Joung and Cheatham Na^+^ ions parameters were added to neutralize the system charge.^47^ In the E·I (reactant) state, we used a bonded model to describe the three zinc finger motifs, where each zinc ion is coordinated with four cysteine residues. This was done by using the ZAFF parameters.^47–50^

In the E-I (product) state, where the Cys26-SSA covalent bond is formed, the zinc finger involved in the inhibition was described with the Y.-P. Pang approach.^49^ In E-I, a water molecule replaces the Cys26 to complete the tetrahedral coordination of the zinc ion^48,50,51^ The custom residue (Figure S15) in which SSA is covalently linked to Cys26 was crafted from the X-ray structure by cutting the Cys26-SSA along the Cα-Cβ bond. A hydrogen atom (H_L_) was added to fill the Cβ vacancy. This moiety was optimized at the DFT-B3LYP/6-311G* level of theory using the Gaussian16 software.^52^ The partial atomic charges were derived using the RESP program to fit the HF/6-31G* electrostatic potentials derived charges using the Merz−Singh−Kollman partitioning scheme. We used the gaff2 force field via the Antechamber (AmberTools23^46^). Conversely, the charges of the backbone atoms of the cysteine moiety were kept restrained to that of the ff14SB force field.

A layer of 14 Å of TIP3P^53^ water molecules was added to all systems, leading to a total system size of ∼320000 atoms. Subsequently, the topologies were converted to the GROMACS format using the parmed software.^54^

#### 4.2.2. Molecular Dynamics Simulations

Classical MD simulations of all the considered systems were done with GROMACS 2022.3 code.^55^ An integration time step of 2 fs was used, and all covalent bonds involving the hydrogen atoms were constrained with the LINCS algorithm.^56^ We used the Particle Mesh Ewald scheme to account for electrostatic interactions,^57^ using a real space cut-off of 10 Å. MD simulations were performed in the isothermal-isobaric NPT ensemble at a temperature of 300 K under the control of the velocity-rescaling thermostat^58^ and the Parrinello-Rahman barostat.^59^ Preliminary energy minimization was performed by employing the steepest descent algorithm. Subsequently, we gradually heated the system to 300 K with an increase of 50 K every 2 ns for a total of 12 ns, keeping the entire system highly restrained except for the solvent atoms and the solute hydrogens. Then we switched to the NPT ensemble, scaling the pressure to 1 bar and using two different barostats: (i) the Berendsen barostat was used for 20 ns with the same restraints on the solute atoms, and (ii) the Parrinello-Rahman barostat for 30 additional ns while leaving the side chains free of restraints. Next, we gradually decreased the restraints in 20 ns. Finally, the reactant and the product were simulated up to 2 μs of MD simulation. Since our investigation started from the E-I deposited structure, we performed two additional replicas of 2 μs each for the E·I state. Indeed, in one of the replicas, a drift of the RMSD was observed in the final part of the simulation for SF3B1 and SF3B3 (Figures S8,S9). The two additional MD simulation replicas of the reactant did not show this trend.

#### 4.2.3. Metadynamics Simulations of SSA in water

2μs-long MD simulation of SSA in pure water was performed before running biased Well-Tempered Metadynamics (WT-MTD) to explore the conformational behavior of spliceostatin A in water solution. This simulation allowed to compute the free energy profile for the rotation of SSA around the dihedral angle connecting the oxygen atom of the tetrahydropyran ring with first carbon atom of the conjugated diene portion (ω_O16,C11,C10,C9,_ Figures S4B, S16), WT-MTD was performed biasing ω_O16,C11,C10,C9_ using GROMACS 2022.3 program patched with PLUMED 2.9.^60^ The errors were calculated by averaging the values of the 3 MTD replicas. The values of ω_O14,C11,C10,C9,_ range from -π to +π, respectively. The parameters employed in WT-MTD simulations were deposition time 5 ps; hill height 0.6 kJ/mol; hill width is π/8 and bias factor 20.

#### 4.2.4. Quantum Mechanics / Molecular Mechanics (QM/MM) MD simulations

Starting from a manually selected frame obtained from the equilibrated part of a 300 ns-long MD simulation of the product state, we performed QM/MM MD simulations with the CP2K v. 11.3 ^61^ program using the same force field parameters of the classical MD simulation for the MM part. Conversely, the QM region was described at the DFT-BLYP-D3 level using the DZVP basis sets to represent the valence orbitals. The electron density was handled using plane waves with a cutoff of 320 Ry. Gaussian-type pseudopotentials following the GTH method were employed to model the atomic cores. Wavefunction optimization utilized the orbital transformation method with a 1 × 10^−5^ convergence threshold for the electronic gradient.

The QM zone, shown in Figure S4, contains 108 or 107 atoms (Lys29^+^ or Lys29) and 7 QM-MM junctions. The QM and MM cut was done along Cα-Cβ of protein residue bond or between two sp^3^-sp^3^ hybridized carbon atoms in SSA. The QM zone includes the side chain of the Asp34 and Lys29 from PHF5A and the three side chains of cysteine residues forming zinc finger 1 (Figures 1C), along with a water molecule. The SSA atoms in the QM region are those up to Carbon atom #10 in Figure 1B and S1. The systems E-I and E×I were relaxed at 300 K by performing 7 and 5.5 ps of QM/MMMD in the NVT ensemble, respectively.

#### 4.2.5. Biased QM/MM MD simulations

The Helmholtz free energy profile along this RC was then investigated via thermodynamic integration by performing constrained QM/MM MD simulation at 9 different RC values (i.e., 1.50, 1.60, 1.65, 1.70, 1.75, 1.80, 2.00, 2.20, and 2.40 Å). The frames of each window were selected from reactive SMD trajectories. The Helmholtz free energy profile was then obtained by integrating the average forces resulting from the QM/MM MD simulations. The standard error for each simulated window was estimated by error propagation analysis from the standard deviation of the constraint force. The overall errors in the activation barrier and reaction-Helmholtz free energy were similarly calculated from the standard error obtained for each window.

#### 4.2.6. Structural and Population Analysis

The root-mean-square deviation (RMSD, Figures S8,S9), root-mean-square fluctuation (RMSF, Figures S10-S12), and the interaction fingerprint that includes hydrogen-bond, hydrophobic, salt-bridge, and π-stacking interactions, were computed with the MDAnalysis^62^ and ProLIF^63^ python libraries. (Table S1)

The free energy profile of the SSA rotation around the dihedral angle ω_O16,C11,C10,C9_ (Figures 1B and S9-S10) was computed using PLUMED 2.9, considering the populations of the relative stated visited in the unbiased MD simulations.^60^ The errors were calculated by averaging the values of the 3 replicas. The energy of SSA rotation in implicit water as solvent was computed at DFT-B3LYP/6-311G* level of theory with a relaxed scan of 12 points in the range -180° to -60 degrees (Figure S3).

The NBO charges were computed through Gaussian 16 software at the DFT-B3LYP/6-311G* level of theory.^52^ The charges were computed on a cluster model in which hydrogen atoms complete the atoms’ valence in the QM zone (Figure S4). The NBO charges were computed on three frames based on the QMMM MD simulation: **A** has the longer (Cys26)S-Zn distance, **B** is the most representative structure of the QM zone, and **C** has the shorted (Cys26)S-Zn distance.

A structural analysis of the Zn-S distances for the zinc fingers present in human spliceosome proteins was done with the biopython package v1.84.^64^ In this analysis of the PDB database, we selected the zinc fingers that (a) possess at least two cysteine residues, (b) and a distorted zinc finger (i.e. their zinc-sulfur distance exceeds 2.4 Å, since this distance is longer than the average of zinc-sulfur bond distances in zinc fingers (Tables S3-S5)).

## 5 Acknowledgments

A. M. and R. R. thank the Italian Association for Cancer Research (AIRC) for financial support (AIRC IG 24514). AM and AP thank “Pathogen Readiness Platform for CERIC-ERIC Upgrade” – PRP@CERIC, financed under the PNRR (National Recovery and Resilience Plan) under Mission 4 “Education and Research” Component 2 “From Research to Enterprise”, Investment Line 3.1 “Fund for the creation of an integrated system of research and innovation infrastructures”, funded by the European Union – Next Generation EU. A.M. and J.A. thanks the PNRR: National Center for Gene Therapy and Drugs based on RNA Technology CUP B83C22002860006 CN_0000004.

The authors thank CINECA, the Italian super-computing center, for computational resources via the “IsB27_EDR” and “IsCc6_SF3bMod” projects.

## Supporting Information

Supplementary data consist of Scheme S1, Supplementary Figures S1-S16, Supplementary Tables S1-S6, and Supplementary Movie 1 (https://doi.org/10.5281/zenodo.15308931).

## Table of Contents Graphics

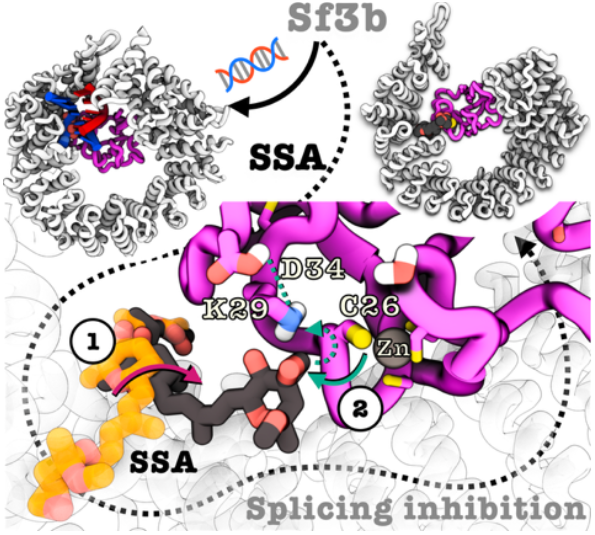

## Notes

https://doi.org/10.5281/zenodo.15308931

### Competing Interest Statement

The authors have declared no competing interest.

### Summary of Updates

Add new figure 3 better explaininginformation already mentioned in the previous version Zinc-fingers in all the figures display now a sulfur-zinc bond and not dashed line Figure 4 and 5 were splitted in two and information already present in the previous version of the text were included in the graphical representation Summary scheme added at the conclusion section. No further calculations /experiments were done respect to the previous version. Part of the text was revised.

