## Supplementary information for "Determinants of spliceostatin reactivity at a spliceosomal zinc finger"

#### Content

- Scheme S1 – General mechanism of covalent inhibition applied to SF3b complex
- Figure S1 – Interaction fingerprint of the reactants
- Figure S2 – SSA atoms involved in the dihedral angle rotation and the dihedral angle values measured in the Classical MD simulation of the reactants
- Figure S3 - QM scan of the SSA dihedral angle
- Figure S4 – Steered MD QM collective variable definition, QM zone used in the QMMM MD simulations, and proton transfer observed in the QMMM MD simulations of the products.
- Figure S5 - Evolution in time of the relevant distances between atoms involved in the transition state
- Figure S6 proton transfer observed in the QMMM MD simulations of the reactants.
- Figure S7 – Interaction fingerprint of the products
- Figure S8 - Root mean square deviation (RMSD) of reactant and product state SF3b complex
- Figure S9 - RMSD of the reactant, other 2 MD replicas
- Figure S10 - Structure of SF3b complex coloured according to the RMSF value of the residues in the reactant and product state
- Figure S11 - Root mean square fluctuation (RMSF) of the reactant and product state SF3B complex
- Figure S12 - RMSF of replicas of the reactant state
- Figure S13 - NBO charges of sulfur atoms of the zinc finger 1
- Figure S14 - Occurrence in the PBD database of elongated Zn-S distances in human spliceosome proteins containing zinc fingers
- Figure S15 – Cys26-SSA non-standard residue definition
- Figure S16 - Well-tempered metadynamics hill's deposition and colvar values

- Table S1 - Interaction fingerprint of SSA with SF3b complex
- Table S2 – propKa Output values for PHF5A-Lys29 and PHF5A-Asp34
- Table S3 - List of the analyzed human proteins that contain zinc fingers
- Table S4 - List of elongated Zn-S distances in the human proteins that contain zinc fingers and are involved in the splicing process
- Table S5 – Occurrence of elongated Zn-S distances in the human proteins that contain zinc fingers and are involved in the splicing process
- Table S6: Acidic and basic side-chain residues in the proximity of distorted zinc-fingers

### Supplementary Schemes

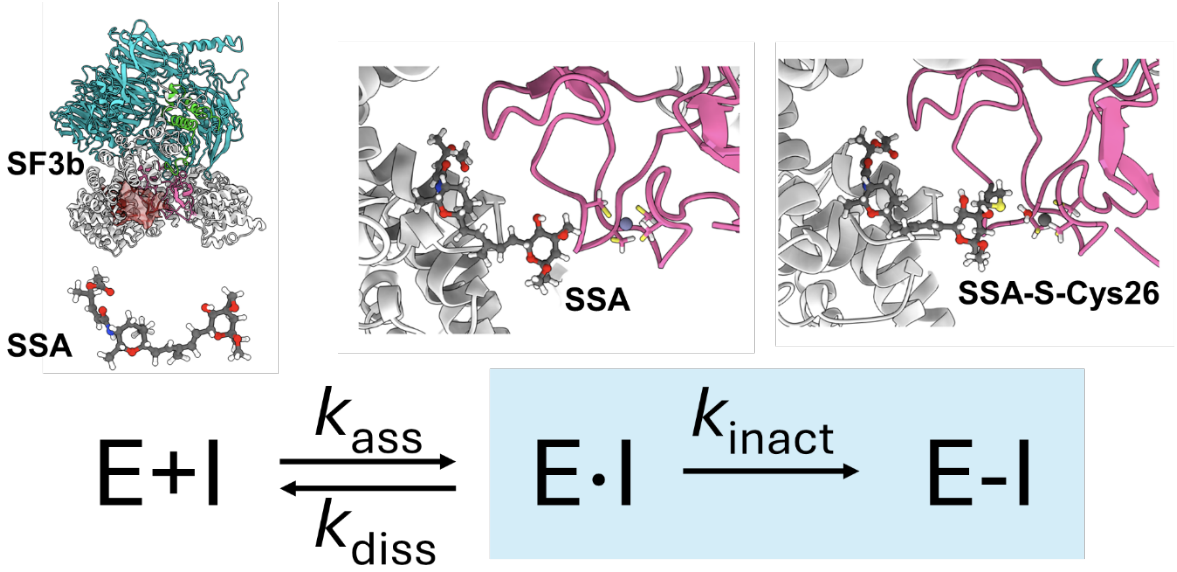


**Scheme S1:** Covalent Inhibition of Spliceostatin A (SSA). Mechanism of covalent inhibition of the SF3b complex (E) by the SSA inhibitor (I). E·I and E–I represent the non-covalent adduct and the covalent complex, respectively, as illustrated in the figures above. The $k_{ass}$, $k_{diss}$, and $k_{inact}$, are the kinetic constants for association, dissociation and inactivation, respectively

### Supplementary Figures

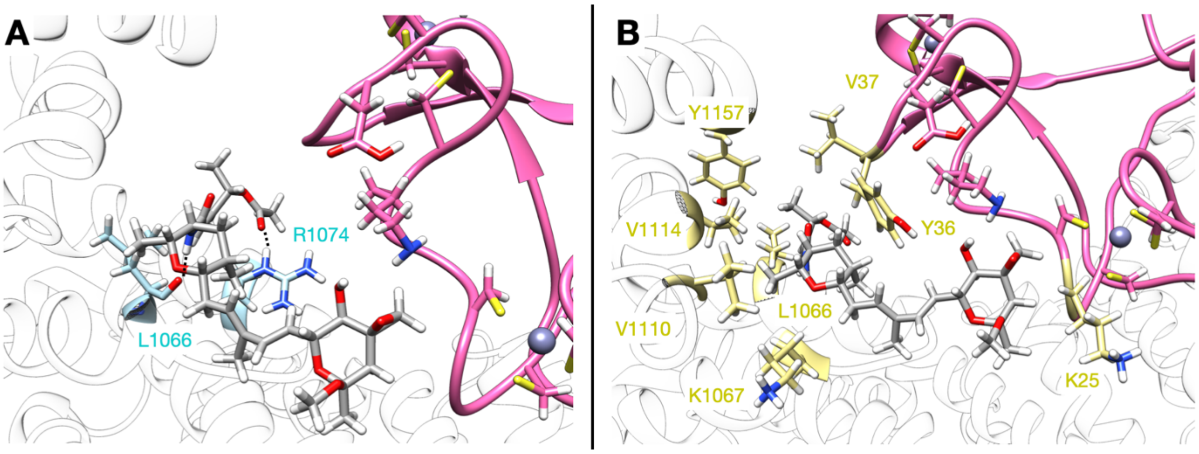


**Figure S1:** Interactions established by SSA with SF3b in the E·I reactant state. SSA carbon atoms are shown in gray. **(A)** Hydrogens bonds are indicated by dashed lines and the carbon atoms of SF3B1 residues involved in hydrogen bonding are colored in light blue. **(B)** Residues establishing the hydrophobic interactions are displayed in pale yellow.

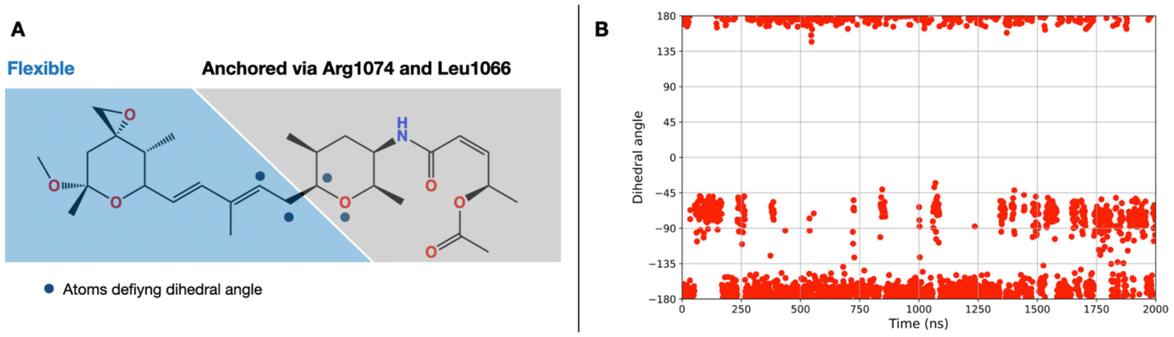


**Figure S2: (A)** SSA structure. The atoms defining the dihedral angle $\omega_{O16,C9,C10,C11}$ for the rotation of SSA are indicated by blue dots. The SSA atoms on the grey background are anchored to SF3B1 protein via Arg1074 and Leu1066, those with the blue background move between the I and L-shape conformation. **(B)** Variation of the dihedral angle during unbiased 2 µs MD simulation.

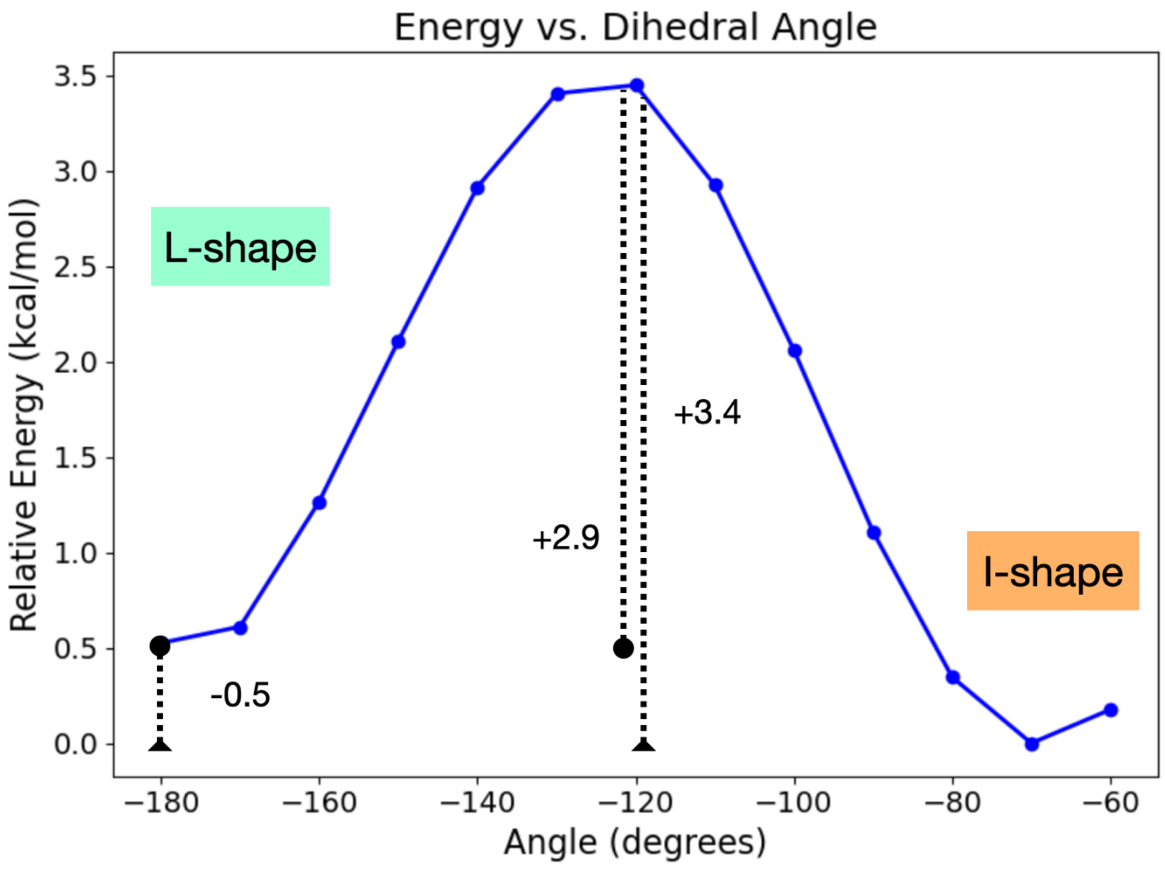


**Figure S3:** Energy for the rotation of SSA along the $\omega_{O16,C9,C10,C11}$ dihedral angle in implicit water as obtained from *DFT-B3LYP* calculations. Energies (kcal/mol) are reported with respect to the I-shape minimum.

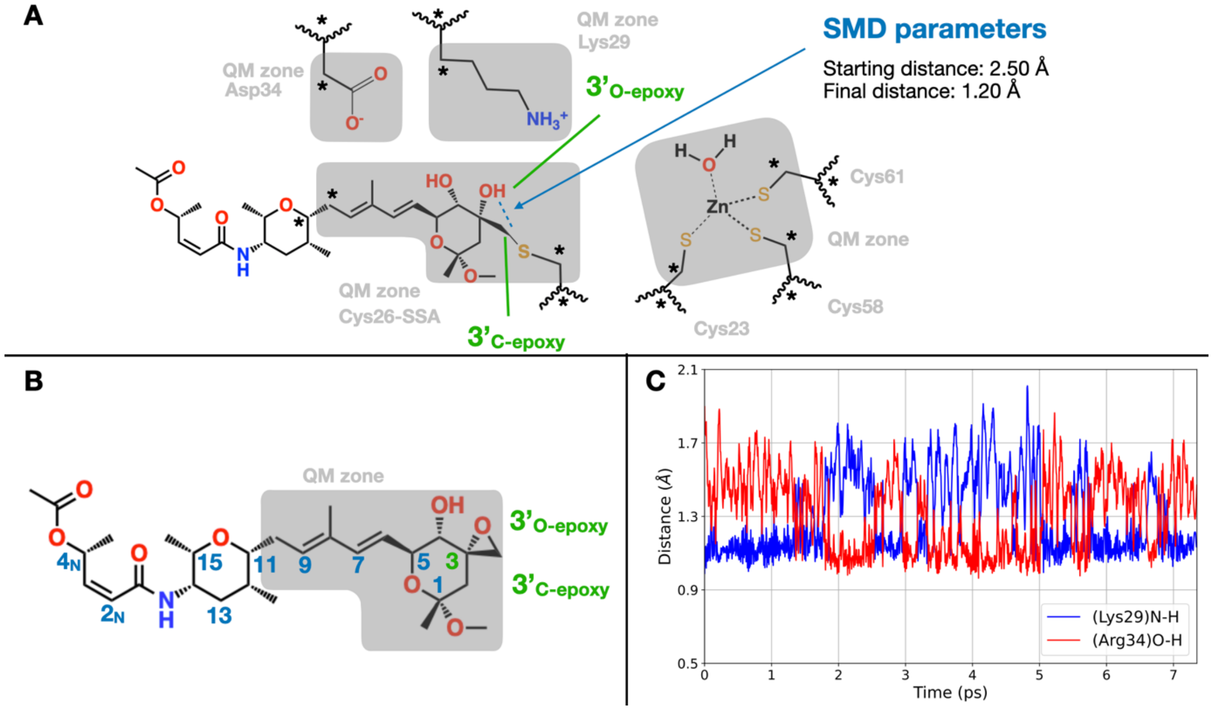


**Figure S4:** The atoms described at quantum mechanical (QM) level in QM/MM molecular dynamics (MD) simulations are shown with grey background. Atoms linking the QM and MM are marked with asterisks, atoms with a white background are in the MM region. **(A)** Representation of the product E-I state QM/MM MD simulation and Steered Molecular Dynamics (SMD) simulation set up, showing the initial and final values of the biased distance. **(B)** Structure of the Spliceostatin A in the reactant state, the atoms of the epoxy warhead are labelled in green. The SSA atoms described at QM level in QM/MM MD simulations are shown with grey background. **(C)** Distance (Å) from Lys29-Nζ (blue) and Asp34-Oδ (red) of the proton shared by $Lys29^{+}$ and $Asp34^{-}$ acid base couple during 7 ps of QM/MM MD simulation of the product E-I state.

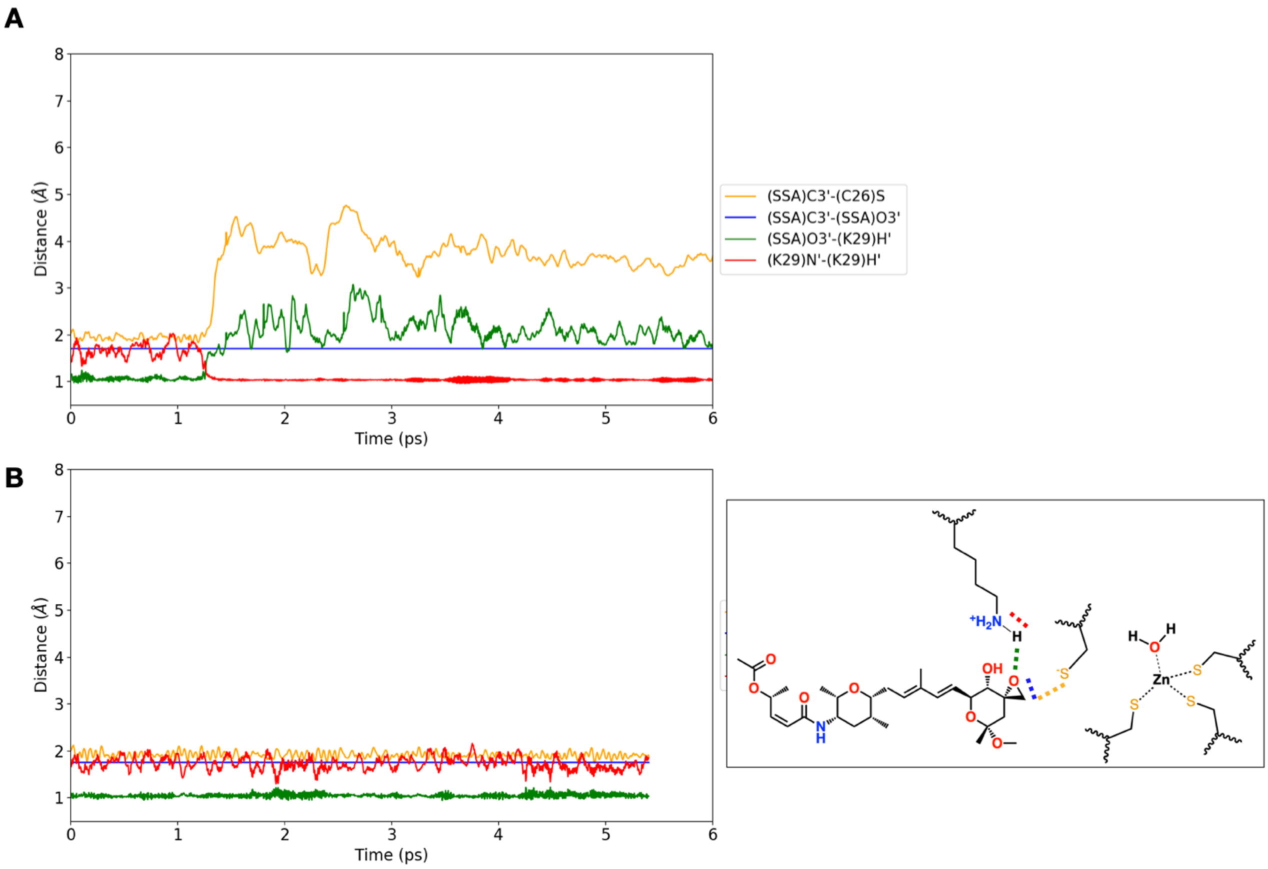


**Figure S5:** Time evolution of the selected distances between reactive atoms at the transition state RC values fixed at 1.70 Å (**A**) and 1.75Å (**B**), show in blue line. The distance between Cys26-S and 3'C atom of SSA is depicted in yellow, the distance between the SSA(O3') and Lys29 hydrogen atom is depicted in green and the distance between the Lys29-Nz and its proton is shown in red. Inset displays a schematic representation of the monitored distances.

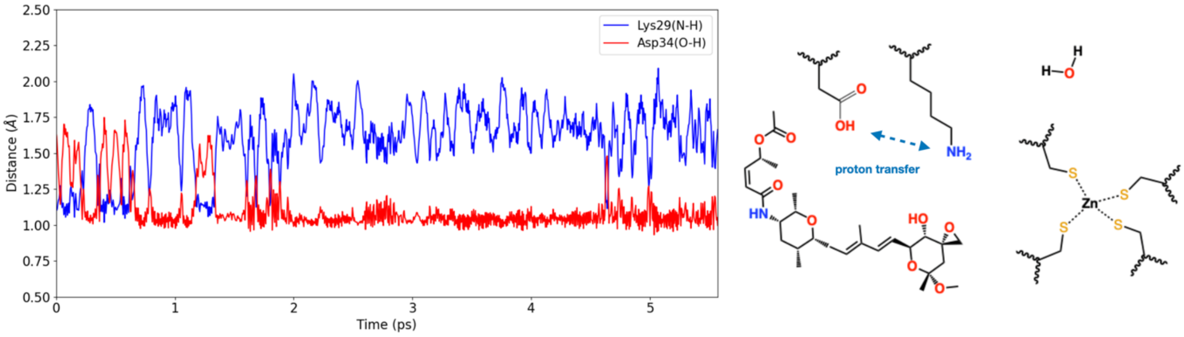


**Figure S6:** Distance (Å) from Lys29-Nz (blue line) and Asp34-Od (red line) of the proton shared by Lys29$^{+}$ and Asp34$^{-}$ acid base couple during 5.5 ps of QM/MM MD simulation of the reactant E·I state. The right panel shows the proton initially present on Lys29-Nζ transfers to Asp34-Oδ during the simulation. The left image depicts the most likely protonation state of the reactant state emerging from QM/MM MD simulations.

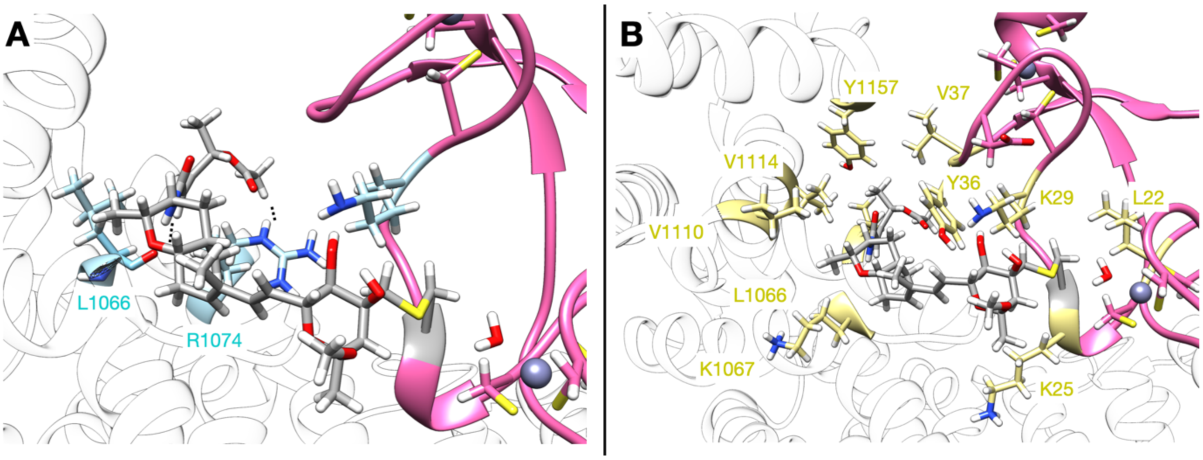


**Figure S7: (A)** Interactions established by SSA with SF3b in the product state. SSA carbon atoms are shown in grey. (A) Hydrogens bonds are indicated by dashed lines and the carbon atoms of SF3B1 residues involved in hydrogen bonding are coloured in light blue. (**B)** Residues establishing the hydrophobic interactions are displayed in pale yellow.

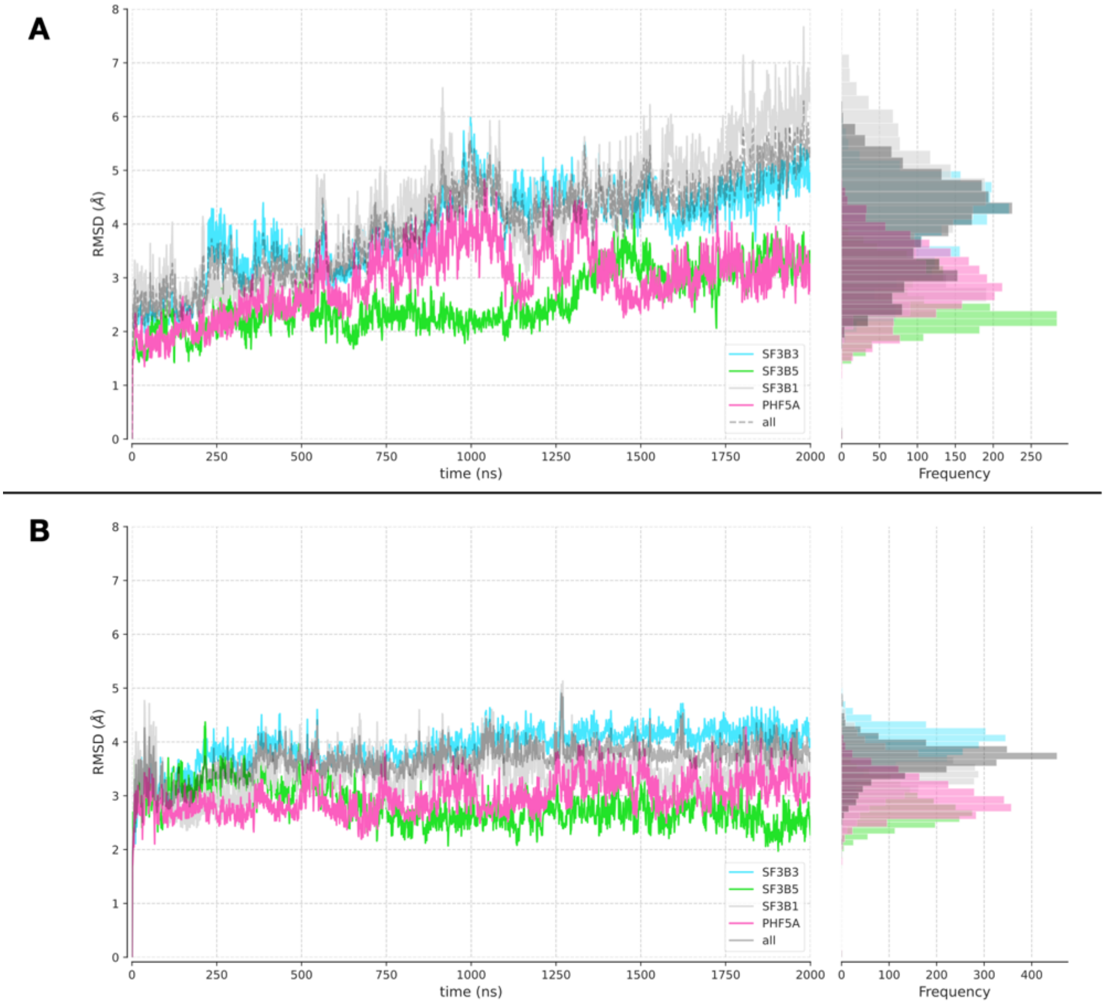


**Figure S8:** Root Mean Square Deviation (RMSD, Å) versus molecular dynamics simulation time (ns) of the SF3b complex components in the non-covalent E·I (reactant) state **(A)**, and in the covalent E-I (product) state **(B)**. The histograms on the side panel shows the distribution of RMSD values.

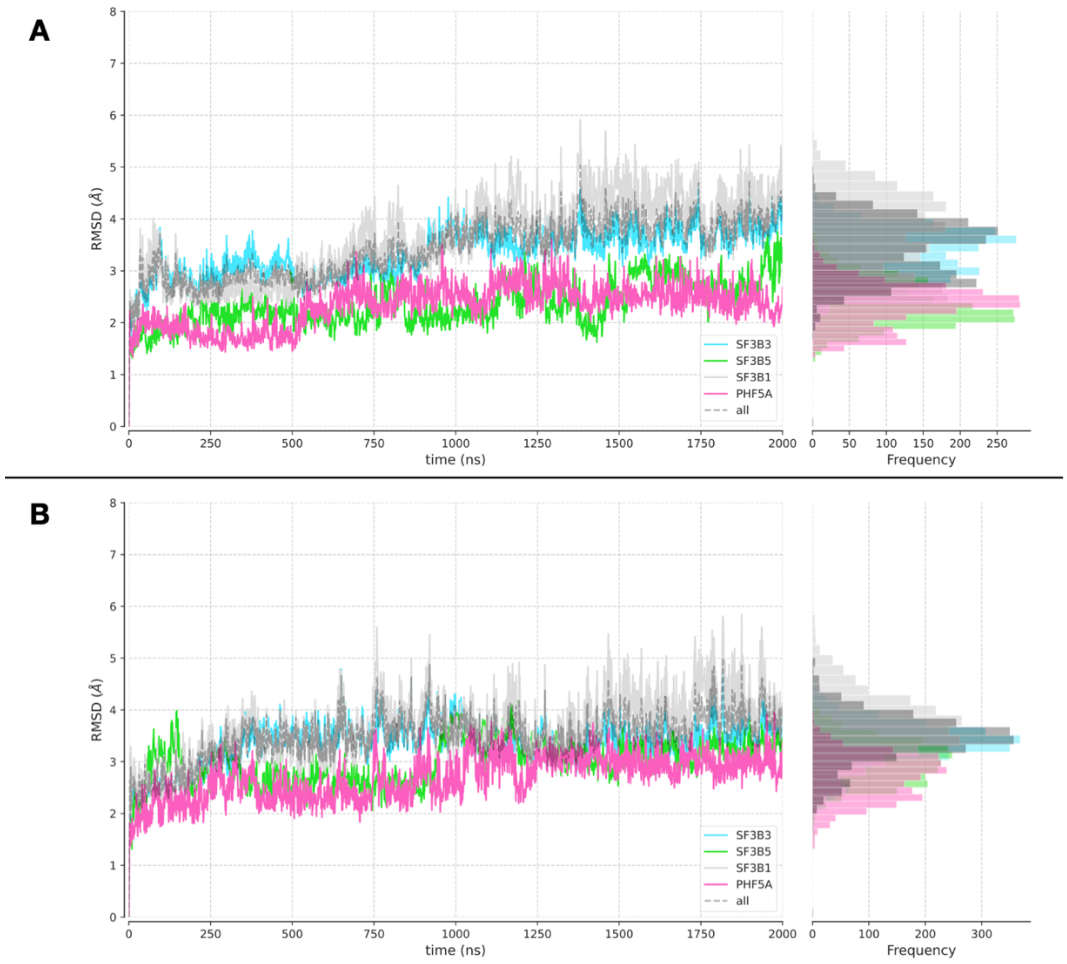


**Figure S9:** Root Mean Square Deviation ((RMSD, Å) versus molecular dynamics simulation time (ns) of the SF3b complex components in the two replicas: replica 1 **(A)**, and replica 2 **(B)** of non-covalent E·I (reactant) state. The histograms on the side panel shows the distribution of RMSD values.

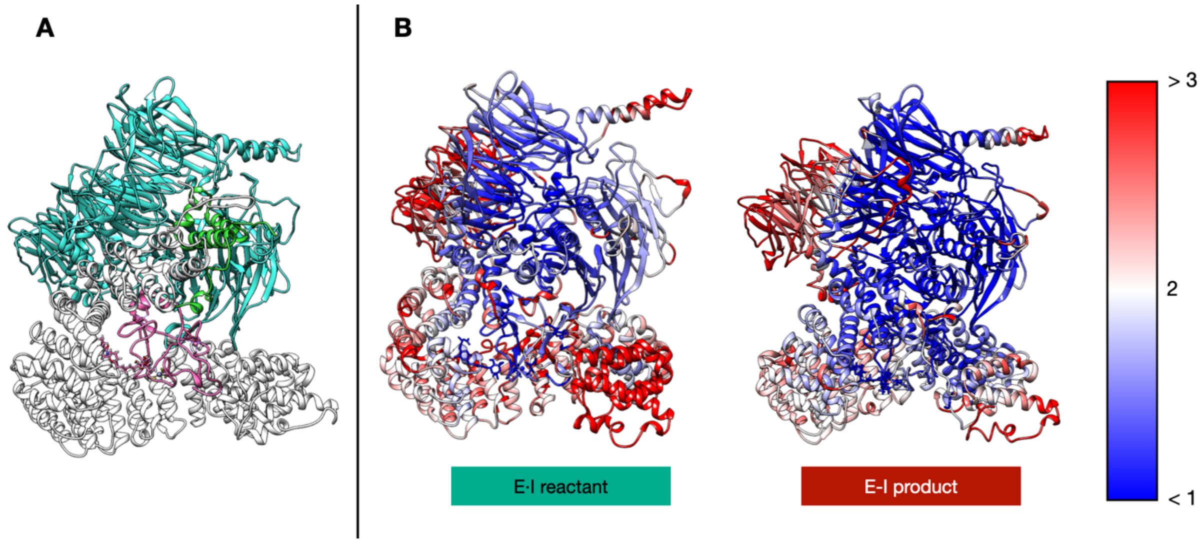


**Figure S10: (A)** Structure of SF3b complex in the E-I state. The SF3B1, SF3B3, SF3B5 and PHF5A proteins are coloured in white, cyan, green, and pink, respectively. **(B)** Structure of SF3b complex coloured according to the RMSF value during the MD simulations in the reactant (E·I) and in the product (E-I) state (center and right respectively).

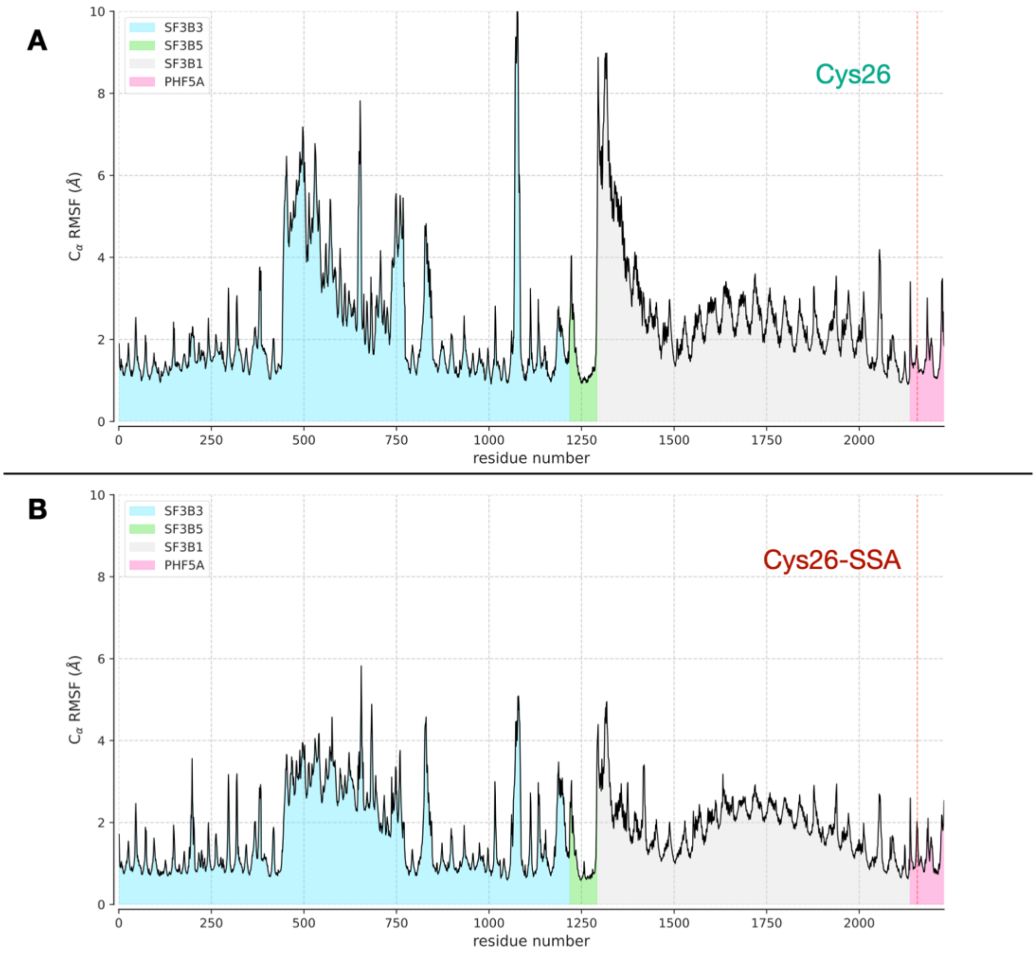


**Figure S11:** Root Mean Square Fluctuation (RMSF, Å) per residue calculated on molecular dynamics simulation trajectories of the SF3b complex in the reactant (**A**) and in the product state (**B**). The areas under the curve are coloured according to the residue location in the SF3b protein components. A red dashed lines marks the position of Cys26 that belongs to a zinc finger in PHF5A in the reactant state and that binds covalently the spliceostatin A in the product state.

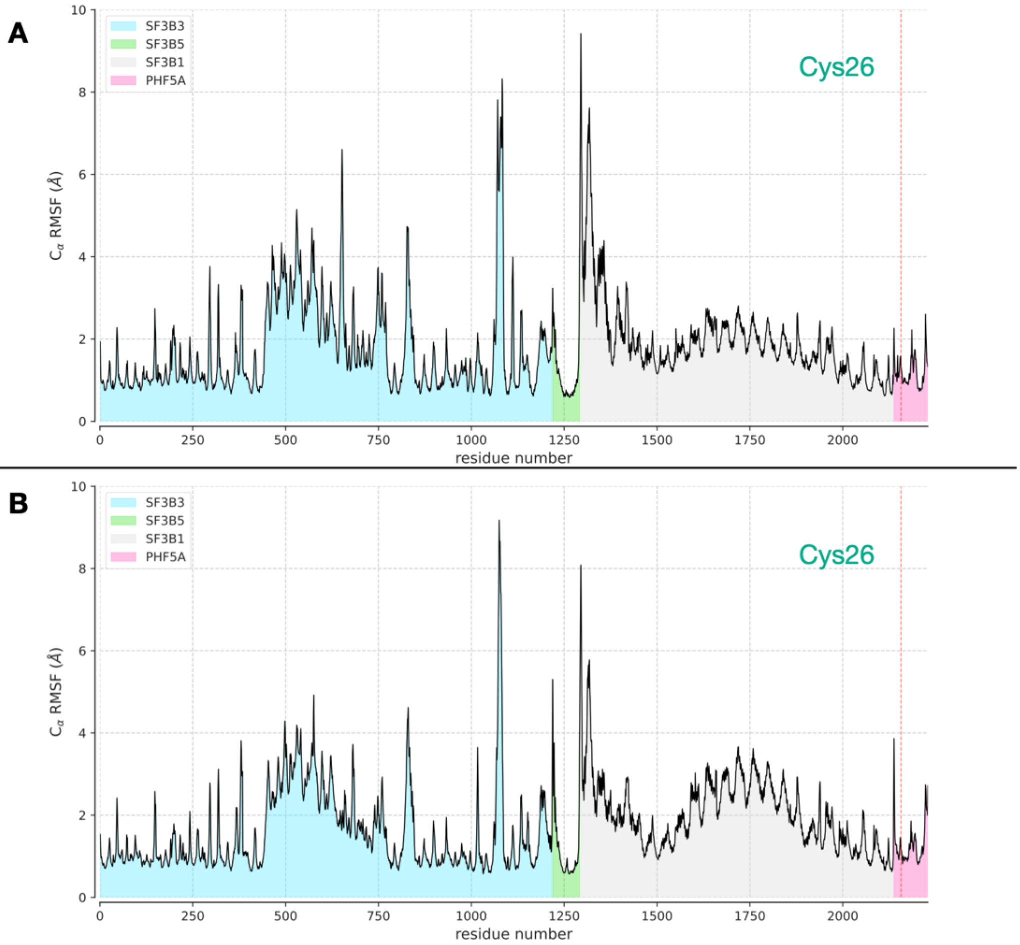


**Figure S12:** Root Mean Square Fluctuation (RMSF, Å) per residue calculated on molecular dynamics simulation trajectories of the SF3b complex in the reactant state of replica 2 and replica 3 in panel (**A**) and (**B**), respectively. The colour scheme is the one reported in Figure S3. A red dashed lines marks the position of Cys26 that belongs to zinc finger 1 in PHF5A in the reactant state.

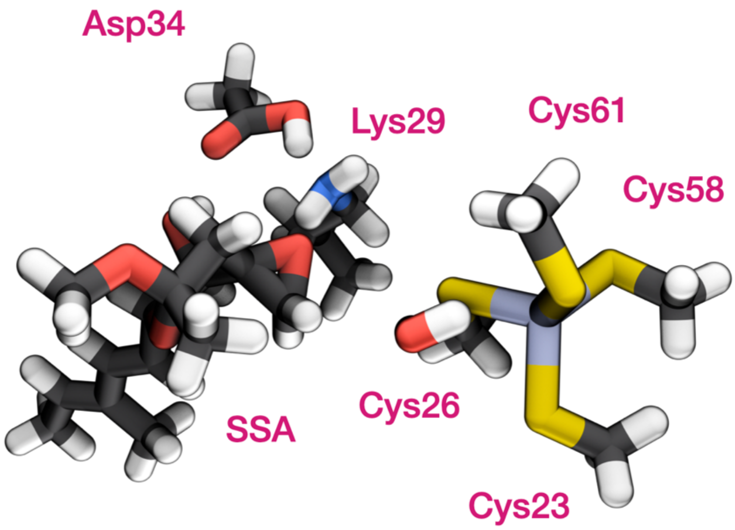


| | Frame A | |
|---|---|---|
| | Distance Zn-S (Å) | Partial charge |
| Cys23 | 2.42 | -0.55 |
| Cys26 | **2.69** | **-0.58** |
| Cys58 | 2.22 | -0.50 |
| Cys61 | 2.50 | -0.59 |
| Zn | - | +0.72 |
| | **Frame B** | |
| | **Distance Zn-S (Å)** | **Partial charge** |
| Cys23 | 2.40 | -0.53 |
| Cys26 | **2.43** | **-0.55** |
| Cys58 | 2.35 | -0.52 |
| Cys61 | 2.40 | -0.55 |
| Zn | - | +0.66 |
| | **Frame C** | |
| | **Distance Zn-S (Å)** | **Partial charge** |
| Cys23 | 2.82 | -0.63 |
| Cys26 | **2.20** | **-0.49** |
| Cys58 | 2.27 | -0.55 |
| Cys61 | 2.36 | -0.55 |
| Zn | - | +0.98 |

**Figure S13**: QM cluster model used for computing the NBO charges. The atoms involved are the one included in the QM zone of the QM/MM molecular dynamics simulation with hydrogen atoms added to the complete the valence of the $sp^3$ carbon atoms. The frame shows the geometry of the QM system in which the Cys26-Zn reaches its maximum value of 2.69 Å. On the right are reported the values obtained for the 3 selected frames. In pink is highlighted the value corresponding to the lowest partial charge and the longest sulfur-zinc distance.

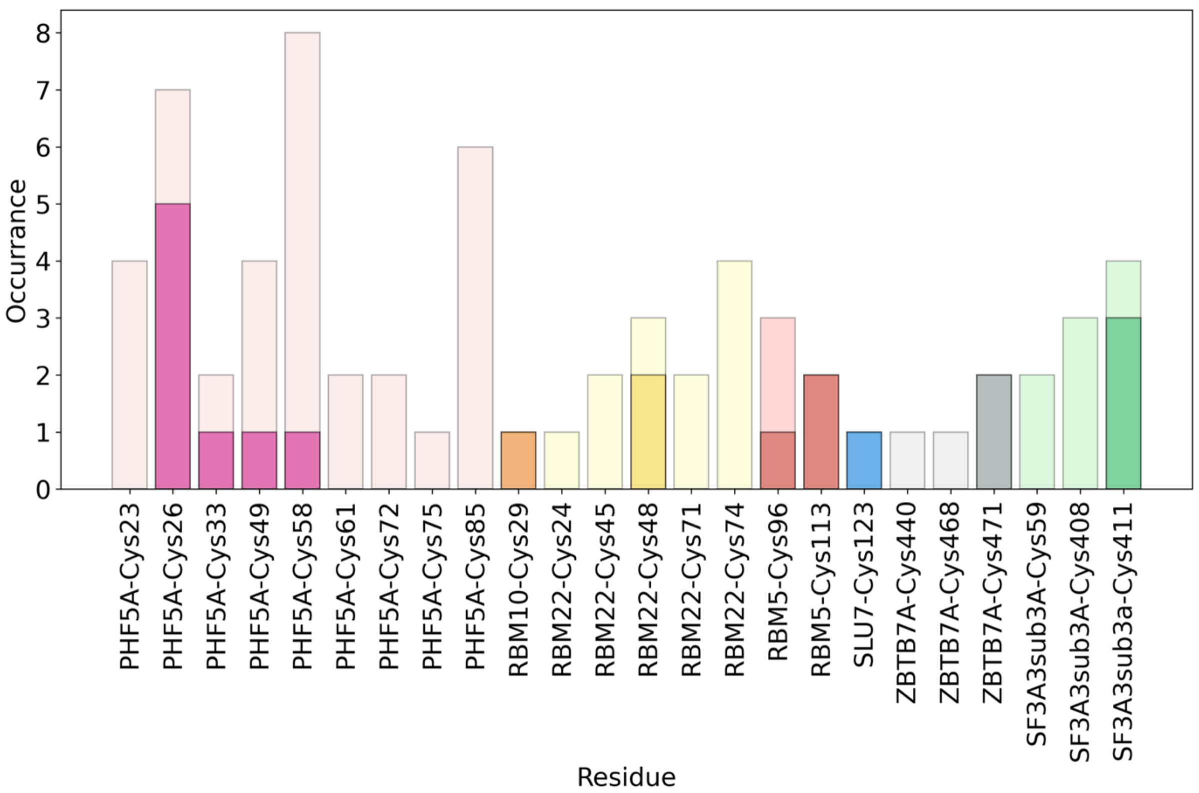

**Figure S14:** Occurrence in the PBD database of elongated Zn-S distances in human spliceosome proteins containing zinc fingers. Light-coloured bars report the occurrence for Zn-S distances above 2.4 Å, dark-coloured bars represent the occurrence of Zn-S distances also exceeding 2.8 Å.

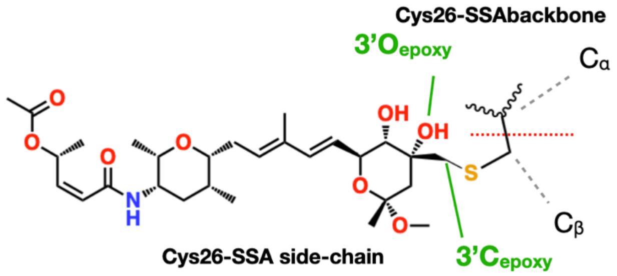


**Figure S15:** Structure of the non-standard residue composed of the Cys26 of PHF5A covalently bound to SSA. The red line delimited the boundaries between the sidechain and the backbone**.**

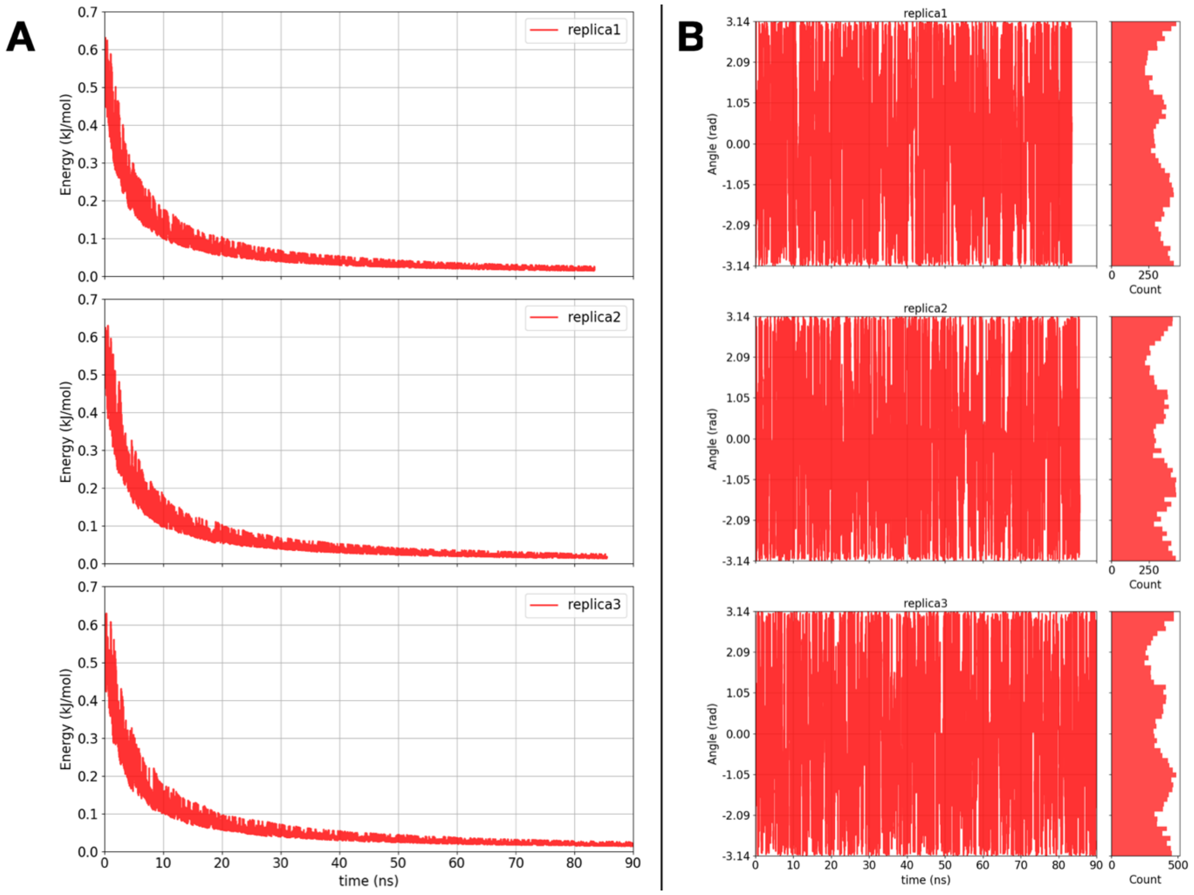

**Figure S16: (A)** Hills height during the WT-MTD of the $\omega_{O16,C9,C10,C11}$ dihedral angle vs simulation time (ns) and **(B)** Actual value of the $\omega_{O16,C9,C10,C11}$ dihedral angle along metadynamics simulation.

### Supplementary Tables

**Table S1:** Interaction fingerprint of SSA with the SF3B1 residues (black, first column) and PHF5A residues (pink, first column) for the Reactant (E·I) (second column) and Product E-I state (third column) during 2 μs of MD. The fourth column reports the occurrence of the interactions of the Product E-I state detected during the 300 ns long MD simulation in with Lys29 was protonated. The last column describes the type of interaction. The last column describes also whether SSA establishes a hydrogen-bond as donor or as acceptor.

| | RS 2μs | PS 2μs | PSp 300ns | Type |
|---|---|---|---|---|
| Tyr36 | 99.7% | 99.7% | 100.0% | Hydrophobic |
| Val1114 | 97.7% | 95.6% | 94.7% | Hydrophobic |
| Val1078 | 97.3% | 97.7% | 99.3% | Hydrophobic |
| Leu1066 | 96.5% | 95.6% | 95.7% | Hydrogen-bond-Donor |
| Leu1066 | | | 95.0% | Hydrophobic |
| Lys1067 | 94.8% | 88.9% | 89.0% | Hydrophobic |
| Arg1074 | 93.1% | 89.5% | 92.4% | Hydrophobic |
| Arg1074 | 90.2% | 80.3% | 92.4% | HB-Acceptor |
| Leu1066 | 89.2% | 90.9% | | Hydrophobic |

| | | | | |
|---|---|---|---|---|
| Val1110 | 78.0% | 60.9% | 85.4% | Hydrophobic |
| Cys23 | | | 66.8% | Hydrophobic |
| Tyr1157 | 64.9% | 63.6% | 69.4% | Hydrophobic |
| Cys1111 | 63.8% | 62.5% | 69.1% | Hydrophobic |
| Val37 | 63.1% | 54.8% | 42.2% | Hydrophobic |
| Lys29 | | 60.2% | 97.3% | Hydrophobic |
| Arg1075 | 58.8% | 50.2% | 63.5% | Hydrophobic |
| His1069 | 58.5% | 55.0% | 70.4% | Hydrophobic |
| Lys25 | | | 56.1% | HB-Donor |
| Lys29 | | 41.4% | | HB-Donor |
| Lys29 | | | 74.8% | HB-Acceptor |
| Leu22 | | 39.6% | 85.4% | Hydrophobic |
| Lys25 | 37.2% | 81.0% | | Hydrophobic |
| Lys29 | | 37.1% | | HB-Acceptor |
| Ile1118 | | 34.6% | 89.4% | Hydrophobic |
| Phe1153 | | | 32.6% | Hydrophobic |

**Table S2:** Predicted pKa with the propKa software[64] for Lys29 and Asp34 belonging to the PHF5A protein.

| | $pK_a$ value |
|---|---|
| Lys29-PHF5A | 9.43 |
| Asp34-PHF5A | 4.68 |

**Table S3:** List of human spliceosome proteins containing zinc fingers as derived from UNIPROT database.

| Protein Name | Gene Name | PDB considered |
|---|---|---|
| E3 ubiquitin-protein ligase RNF113A (EC 2.3.2.27) (Cwc24 homolog) (RING finger protein 113A) (Zinc finger protein 183) | RNF113A RNF113 ZNF183 | 5Z56;5Z58;6FF4;6FF7;7DVQ;7QTT;8CH6; |
| Zinc finger and BTB domain-containing protein 7A (Factor binding IST protein 1) (FBI-1) (Factor that binds to inducer of short transcripts protein 1) (HIV-1 1st-binding protein 1) (Leukemia/lymphoma-related factor) (POZ and Krueppel erythroid myeloid ontogenic factor) (POK erythroid myeloid ontogenic factor) (Pokemon) (Pokemon 1) (TTF-I-interacting peptide 21) (TIP21) (Zinc finger protein 857A) | ZBTB7A FBI1 LRF ZBTB7 ZNF857A | 2IF5;2NN2;7EYI;7N5S;7N5T;7N5U;7N5V;7N5W; 8E3D;8E3E;8H9H; |
| Pre-mRNA-splicing factor SLU7 (hSlu7) | SLU7 | 5XJC;6ICZ;6QDV;7W5A;7W5B;8C6J; |
| U1 small nuclear ribonucleoprotein C (U1 snRNP C) (U1-C) (U1C) | SNRPC | 2VRD;3CW1;4PJO;6ELD;6QX9;7VPX; |
| RNA-binding protein 5 (Protein G15) (Putative tumor suppressor LUCA15) (RNA-binding motif protein 5) (Renal carcinoma antigen NY-REN-9) | RBM5 H37 LUCA15 | 2LK0;2LK1;2LKZ;5MF9;5MFY;7PCV;7PDV; |

| | | |
|---|---|---|
| RNA-binding protein 10 (G patch domain-containing protein 9) (RNA-binding motif protein 10) (RNA-binding protein S1-1) (S1-1) | RBM10 DXS8237E GPATC9 GPATCH9 KIAA0122 | 2LXI;2M2B;2MXV;2MXW;5ZSW;5ZSY; |
| Splicing factor 3A subunit 3 (SF3a60) (Spliceosome-associated protein 61) (SAP 61) | SF3A3 SAP61 | 2DT7;5Z56;5Z57;5Z58;6AH0;6AHD;6FF7; 6QX9;6Y50;6Y53;6Y5Q;7ABG;7ABH;7ABI; 7EVO;7ONB;7Q3L;7Q4O;7Q4P;7QTT;7VPX; 8CH6;8H6E;8H6J;8H6K;8H6L;8HK1;8QO9; 8QXD;8QZS;8R08;8R09;8R0A;8R0B;8RM5; |
| Splicing factor 1 (Mammalian branch point-binding protein) (BBP) (mBBP) (Transcription factor ZFM1) (Zinc finger gene in MEN1 locus) (Zinc finger protein 162) | SF1 ZFM1 ZNF162 | 1K1G;1O0P;1OPI;2M09;2M0G;4FXW;4FXX; 7VH9;7VPX;8PXX; |
| PHD finger-like domain-containing protein 5A (PHD finger-like domain protein 5A) (Splicing factor 3B-associated 14 kDa protein) (SF3b14b) | PHF5A | 5IFE;5O9Z;5SYB;5Z56;5Z57;5Z58;5ZYA;6AH0; 6AHD;6EN4;6FF4;6FF7;6QX9;6Y50;6Y5Q;7ABG; 7ABH;7ABI;7B0I;7B91;7B92;7B9C;7DVQ;7EVN;7 EVO;7OMF;7ONB;7OPI;7Q3L;7Q4O;7Q4P;7QTT;7 VPX;8CH6;8H6E;8H6J;8H6K;8H6L;8HK1;8QO9; 8QXD;8QZS;8R08;8R09;8R0A;8R0B;8RM5;8Y7E; |
| Pre-mRNA-splicing factor RBM22 (RNA-binding motif protein 22) (Zinc finger CCCH domain-containing protein 16) | RBM22 ZC3H16 199G4 | 2YTC;5MQF;5XJC;5YZG;5Z56;5Z57;6FF4;6FF7;6ICZ; 6ID0;6ID1;6QDV;6ZYM;7A5P;7AAV;7ABG;7ABI;7QTT; 7W59;7W5A;7W5B;8C6J;8CH6; |
| Pre-mRNA-processing factor 19 (EC 2.3.2.27) (Nuclear matrix protein 200) (PRP19/PSO4 homolog) (hPso4) (RING-type E3 ubiquitin transferase PRP19) (Senescence evasion factor) | PRPF19 NMP200 PRP19 SNEV | 4LG8;5MQF;5XJC;5YZG;5Z56;5Z57;6FF7;6ICZ;6ID0; 6ID1;6QDV;7A5P;7W59;7W5A;7W5B;8C6J;8CH6; |
| Peptidyl-prolyl cis-trans isomerase E (PPIase E) (EC 5.2.1.8) (Cyclophilin E) (Cyclophilin-33) (Rotamase E) | PPIE CYP33 | 1ZMF;2CQB;2KU7;2KYX;2R99;3LPY;3MDF;3UCH; 5MQF;5YZG;5Z56;5Z57;6FF7;6ICZ;6ID0;6ID1;7A5P; 7ABI;7W59;7W5A;7W5B;7ZEV;7ZEW;7ZEX;7ZEY;7ZEZ; 8C6J;8CH6; |
| Serine/arginine repetitive matrix protein 2 (300 kDa nuclear matrix antigen) (Serine/arginine-rich splicing factor-related nuclear matrix protein of 300 kDa) (SR-related nuclear matrix protein of 300 kDa) (Ser/Arg-related nuclear matrix protein of 300 kDa) (Splicing coactivator subunit SRm300) (Tax-responsive enhancer element-binding protein 803) (TaxREB803) | SRRM2 KIAA0324 SRL300 SRM300 HSPC075 | 5MQF;5XJC;5YZG;5Z56;5Z57;6FF4;6FF7;6ICZ;6QDV; 6ZYM;7A5P;7DVQ;7QTT;7W59;7W5A;7W5B;8C6J;8CH6; |

**Table S4:** List of elongated Zn-S distances in the human spliceosome proteins that contain zinc finger (ZF) motifs. The distances reported are those exceeding 2.40 Å. The first column reports the PDB code, the second the experimental method used to solve the structure, the third the resolution, the fourth the protein name, the fifth the type of zinc finger motif, the fourth the elongated Cys residue and the actual value of the distance.

| PDB | Method | Resolution | Protein name | ZF type | residue |
|---|---|---|---|---|---|
| 2MXV | NMR | | RBM10 | Cys4 | Cys-29: 2.84 |
| 4PJO | X-Ray diffraction | 3.30 Å | U1 snRNP C | 2Cys2His | Cys-6 (Chain l): 2.42 |
| 5IFE | X-Ray diffraction | 3.10 Å | PHF5A | Cys4 | Cys-26 (Chain D): 2.76 |
| | | | | Cys4 | Cys-75 (Chain D): 2.44 |
| 5ZYA | Electron Microscopy | 3.95 Å | PHF5A | Cys4 | Cys-23 (Chain D): 2.49 |
| | | | | Cys4 | Cys-26 (Chain D): 3.10 |
| | | | | Cys4 | Cys-58 (Chain D): 2.50 |
| | | | | Cys4 | Cys-61 (Chain D): 2.48 |
| | | | | Cys4 | Cys-11 (Chain D): 2.56 |
| | | | | Cys4 | Cys-46 (Chain D): 2.47 |
| | | | | Cys4 | Cys-49 (Chain D): 3.13 |
| | | | | Cys4 | Cys-85 (Chain D): 2.53 |
| 6EN4 | X-Ray diffraction | 3.08 Å | PHF5A | Cys4 | Cys-23 (Chain D): 2.46 |
| | | | | Cys4 | Cys-26 (Chain D): 3.06 |
| | | | | Cys4 | Cys-58 (Chain D): 2.41 |
| | | | | Cys4 | Cys-61 (Chain D): 2.43 |
| | | | | Cys4 | Cys-85 (Chain D): 2.58 |

| | | | | | |
|---|---|---|---|---|---|
| 6FF4 | Electron Microscopy | 3.40 Å | RBM22 | 4Cys | Cys-74 (Chain P): 2.55 |
| | | | PHF5A | 4Cys | Cys-23 (Chain y): 2.42 |
| | | | | Cys4 | Cys-58 (Chain y): 2.73 |
| | | | | Cys4 | Cys-30 (Chain y): 2.50 |
| | | | | Cys4 | Cys-33 (Chain y): 3.45 |
| | | | | Cys4 | Cys-72 (Chain y): 2.51 |
| 6ICZ | Electron Microscopy | 3.0 Å | RBM22 | Cys4 | Cys-48 (Chain O): 2.57 |
| 6ID0 | Electron Microscopy | 2.9 Å | RBM22 | Cys4 | Cys-45 (Chain O): 2.50 |
| | | | | Cys4 | Cys-48 (Chain O): 3.88 |
| | | | | Cys4 | Cys-71 (Chain O): 2.52 |
| | | | | Cys4 | Cys-74 (Chain O): 2.52 |
| 6ID1 | Electron Microscopy | 2.86 Å | RBM22 | Cys4 | Cys-45 (Chain O): 2.56 |
| | | | | Cys4 | Cys-48 (Chain O): 3.93 |
| | | | | Cys4 | Cys-71 (Chain O): 2.65 |
| | | | | Cys4 | Cys-74 (Chain O): 2.52 |
| 6Y50 | Electron Microscopy | 4.10 | PHF5A | Cys4 | Cys-11 (Chain y): 3.00 |
| 7B92 | X-Ray diffraction | 3.0 Å | PHF5A | Cys4 | Cys-85 (Chain D): 2.74 |
| | | | | Cys4 | Cys-30 (Chain D): 2.47 |
| | | | | Cys4 | Cys-58 (Chain D): 2.68 |
| 7B9C | X-Ray diffraction | 3.4 Å | PHF5A | Cys4 | Cys-11 (Chain D): 2.42 |
| 7EVN | Electron Microscopy | 2.57 Å | PHF5A | Cys4 | Cys-49 (Chain D): 2.57 |
| 7EVO | Electron Microscopy | 2.5 Å | PHF5A | Cys4 | Cys-58 (Chain 6): 2.64 |
| | | | | Cys4 | Cys-49 (Chain 6): 2.61 |
| | | | SF3A3 sub 3A | 2Cys2his | Cys-408 (Chain C): 2.47 |
| | | | SF3A3 sub 3A | 2Cys2his | Cys-411 (Chain C): 3.37 |
| 7EYI | X-Ray diffraction | 2.4 Å | zinc-finger and BTB domain-containing protein 7A | 3Cys1his | Cys-468 (Chain G): 2.51 |
| | | | | 3Cys1his | Cys-471 (Chain G): 2.90 |
| | | | | 3Cys1his | Cys-440 (Chain H): 2.58 |
| | | | | 3Cys1his | Cys-471 (Chain H): 3.20 |
| | | | | 3Cys1his | Cys-490 (Chain H): 2.48 |
| 7ONB | Electron Microscopy | 3.10 Å | PHF5A | Cys4 | Cys-26 (Chain D): 3.80 |
| 7PDV | X-Ray diffraction | 3.49 Å | RBM5 | Cys3* | Cys-96 (Chain A): 2.72 |
| | | | RBM5 | Cys4 | Cys-96 (Chain E): 2.47 |
| | | | RBM5 | Cys4 | Cys-113 (Chain E): 3.46 |
| | | | RBM5 | Cys4 | Cys-96 (Chain G): 2.96 |
| | | | | Cys4 | Cys-113 (Chain G): 3.06 |
| 7Q3L | Electron Microscopy | 2.21 Å | PHF5A | Cys4 | Cys-26 (Chain G): 2.80 |
| 7Q4O | Electron Microscopy | 2.21 Å | SF3A3 sub 3A | 2Cys2His | Cys-411 (Chain 9): 2.78 |
| | | | PHF5A | Cys4 | Cys-11 (Chain G): 2.52 |
| | | | | Cys4 | Cys-85 (Chain G): 2.55 |
| | | | | Cys4 | Cys-26 (Chain G): 2.43 |
| | | | | Cys4 | Cys-58 (Chain G): 2.67 |
| 7Q4P | Electron Microscopy | 2.15 Å | SF3A3 sub 3A | 2CysHhis | Cys-59 (Chain 1): 2.55 |
| | | | PHF5A | Cys4 | Cys-30 (Chain G): 2.46 |
| | | | | Cys4 | Cys-11 (Chain G): 2.49 |

| | | | | | |
|---|---|---|---|---|---|
| | | | | Cys4 | Cys-85 (Chain G): 2.61 |
| | | | | Cys4 | Cys-26 (Chain G): 2.87 |
| | | | | Cys4 | Cys-58 (Chain G): 2.89 |
| 7VPX | Electron Microscopy | 3.0 Å | SF3A3 sub 3A | 2Cys2His | Cys-59 (Chain B): 2.49 |
| | | | SF3A3 sub 3A | 2Cys2His | Cys-408 (Chain C): 2.47 |
| | | | | 2Cys2His | Cys-411 (Chain C): 3.37 |
| | | | PHF5A | Cys4 | Cys-85 (Chain 6): 2.78 |
| | | | | Cys4 | Cys-33 (Chain 6): 2.62 |
| | | | | Cys4 | Cys-72 (Chain 6): 2.68 |
| | | | | Cys4 | Cys-23 (Chain 6): 2.55 |
| 7W59 | Electron Microscopy | 3.6 Å | RBM22 | Cys4 | Cys-24 (Chain O): 2.63 |
| | | | | Cys4 | Cys-74 (Chain O): 2.58 |
| 7W5A | Electron Microscopy | 3.6 Å | SLU7 | Cys3His1 | Cys-123 (Chain 1): 3.40 |
| 8HK1 | Electron Microscopy | 3.6 Å | PHF5A | Cys4 | Cys-58 (Chain 6): 2.64 |
| | | | | Cys4 | Cys-49 (Chain 6): 2.61 |
| | | | SF3A3 sub 3A | 2Cys2His | Cys-408 (Chain C): 2.47 |
| | | | | 2Cys2His | Cys-411 (Chain C): 3.37 |

**Table S5:** List of elongated Zn-S distances in the human spliceosome proteins that contain zinc finger motifs. The distances reported are the one that exceed 2.5 Å. The first column reports the residues that show an elongated Zn-S distance, the second the number of occurrences in the PDBs examined in Table S3, and column 3 reports in how many PDBs that distance exceeds 2.8 Å. The percentages of occurrence respect to the number of analysed PDB in which the protein is present is reported in parenthesis.

| Protein and residue | Occurrence | Occurrence above 2.8Å |
|---|---|---|
| PHF5A-Cys-23 | 4 (8.3%) | 0 |
| PHF5A-Cys-26 | 7(14.6%) | 5 (10.4%) |
| PHF5A-Cys-33 | 2 (4.2%) | 2(4.2%) |
| PHF5A-Cys-49 | 4 (8.3%) | 1 (2.1%) |
| PHF5A-Cys-58 | 8 (16.7%) | 1 (2.1%) |
| PHF5A-Cys-61 | 2 (4.2%) | 0 |
| PHF5A-Cys-72 | 2 (4.2%) | 0 |
| PHF5A-Cys-75 | 1 (2.1%) | 0 |
| PHF5A-Cys-85 | 6 (12.5%) | 0 |
| RBM10-Cys-29 | 1 (16.7%) | 1 (16.7%) |
| RBM22-Cys-24 | 1 (4.3%) | 0 |
| RBM22-Cys-45 | 2 (8.7%) | 0 |
| RBM22-Cys-48 | 3 (13.0%) | 2 (8.7%) |
| RBM22-Cys-71 | 2 (8.7%) | 0 |
| RBM22-Cys-74 | 4 (17.4%) | 0 |

| | | |
|---|---|---|
| RBM5-Cys-96 | 3 (42.3%) | 1 (14.3%) |
| RBM5-Cys-113 | 2 (28.3%) | 2 (28.3%) |
| SLU7-Cys-123 | 1 (16.7%) | 1 (16.7%) |
| ZBTB7A-Cys-440 | 1 (9.1%) | 0 |
| ZBTB7A-Cys-468 | 1 (9.1%) | 0 |
| ZBTB7A-Cys-471 | 2 (18.2%) | 2 (18.2%) |
| SF3A3 sub 3a-Cys-59 | 2 (5.7%) | 0 |
| SF3A3 sub 3a-Cys-408 | 3 (8.6%) | 0 |
| SF3A3 sub 3a-Cys-411 | 4 (11.4%) | 3 (8.6%) |

**Table S6:** Cysteine showing an elongated Zn-S bond (according to Table S5) are reported in the first column. The second column lists the other residues constituent of the zinc finger. Third and fourth column shows acidic and basic residues in the proximity of distorted zinc fingers. Where it is not specified the residues belong to the same chain (details in Table S4). A dash indicates that no acidic side-chain residue was detected. Similarly to what observed in the E-I state, we considered the side chain of the basic residues within 5 Å from the distorted zinc-finger and the side-chain of acidic residues are within 5 Å from the basic residues.

| | **Other residues in the ZF** | **Basic side chain residue** | **Acid side chain residue** |
|---|---|---|---|
| RBM10-Cys29 | Cys26, Cys12, Cys15 | Lys14 | Glu35 |
| RBM22-Cys45 | Cys48, Cys71, Cys74 | Lys46 | Glu69 |
| RBM22-Cys48 | Cys45, Cys71, Cys74 | Arg50 | Glu44 |
| RBM22-Cys48 | Cys45, Cys71, Cys74 | Arg50 | Glu122 |
| RBM5-Cys96 chain G | Cys110, Cys113, Cys99 | Lys98 | - |
| RBM5-Cys113 chain G * | Cys110, Cys96, Cys99 | Arg112 | - |
| RBM5-Cys113 chain E | Cys110, Cys96, Cys99 | Lys98 | His58 chain A |
| SLU7-Cys123 | Cys120, His128, Cys133 | Arg138 | Glu135 |
| ZBTB7A-Cys471 | Cys468, His484, Cys490 | Lys473 | - |
| ZBTB7A-Cys471 | Cys468, His484, Cys490 | His212 | - |
| SF3A3 sub 3a-Cys411 | Cys408, His425, His431 | Arg340 | His433 |
| SF3A3 sub 3a-Cys411 | Cys408, His425, His431 | Arg340 | Glu428 |
